# GMP-Cor: Quantifying Transcriptional Dysregulation in Antibiotic Tolerance and Persistence

**DOI:** 10.64898/2026.11.02.763672

**Authors:** Alon Gutfreund, Adi Rotem, Irine Ronin, Adam Z. Rosenthal, Oded Agam, Nathalie Q. Balaban

## Abstract

When cells are exposed to stress, they typically activate stress-response pathways and enter a regulated growth-arrest. But what if the stress overwhelms these pathways? In such cases, cells may enter a *disrupted growth-arrest* - a poorly adapted, abrupt cessation of growth, characterized by a broad distribution of lag times upon stress removal [1]. This distribution underlies antibiotic persistence during stress recovery[2]. Therefore, gaining a deeper understanding of the disrupted growth-arrest is important for developing more effective antibiotic therapies. In this study, we perform single cell RNA-seq on bacteria in disrupted conditions. We develop GMP-Cor: a metric to quantify dysregulation and analyze gene-gene correlation patterns in sparse single-cell data. We find that gene-gene correlations are significantly reduced in disrupted cells compared to regulated cells, providing direct evidence that this tolerance-enhanced state, along with its increased membrane sensitivity, is associated with global-dysregulation. We demonstrate that GMP-Cor can extend to other biological systems, such as persistent cancer cells, where it separates proliferating from drug-arrested cells[3].

## 1 Introduction

A cell functions as a complex machine, refined during evolution, with its gene regulatory network governing the activation or suppression of specific pathways. This network coordinates interactions between genes, whose collective behavior controls cellular homeostasis. In stressful environments, specific genetic pathways are activated as part of a regulated stress response. For example, when an *E. coli* culture gradually exhausts nutrients, it enters stationary phase through a well-characterized regulatory program, controlled by the alternative sigma factor RpoS (σ*^S^*) which redirects transcription toward stationary-phase and general stress-response genes[4, 5, 6, 7, 8, 9, 10]. Together with the alarmone (p)ppGpp [11], this produces a reversible, growth-arrested state. During such a regulated growth-arrest, bacteria are more protected from various forms of stress [12, 13] and highly tolerant to antibiotic treatments [14, 15, 16]. Upon nutrient replenishment, bacteria resume growth promptly and tolerance is lost [17, 18]. Crucially, such stress response regulatory programs require adequate time to respond and high coordination.

While growth-arrest in mammalian cells is finely categorized into distinct physiological states - such as quiescence, senescence, or apoptosis - growth-arrested bacteria are traditionally described under the broad term “dormant”. However, bacteria that arrest their growth may be in very different physiologies [19, 13, 20]. A classification has recently emerged that draws a fundamental distinction between two archetypes of growth arrest: regulated (Reg-Arrest) and disrupted (Dis-Arrest)[2]. Reg-Arrest is a controlled entry into stationary phase, allowing the cell to activate mechanisms for prolonged survival under stress. Dis-Arrest, in contrast, occurs when an acute stress overwhelms the regulatory network, resulting in reduced adaptability, and uncontrolled entry into a distinct growth-arrested state[1, 2]. The added value of the Dis-Arrest framework is a set of quantitative predictions that do not follow from the absence of regulation alone. It establishes how to distinguish between Reg-Arrest and Dis-Arrest without knowing the history of treatment. It predicts that the duration of the arrest shapes the distribution of lag times [1]; that disruption confers antibiotic tolerance while creating collateral sensitivity to membrane-targeting agents [2]; and - the subject of the present work - that gene-gene correlations and coordination of pathways are reduced.

Acute stress leading to Dis-Arrest can arise from a range of conditions such as exposure to chemicals, unnatural starvation, or the immune response of the host, each targeting distinct pathways [21, 1, 22, 23]. As cells do not grow in this state, they typically exhibit antibiotic tolerance, enabling them to withstand treatments that target only actively growing bacteria [14, 24, 25, 26]. When the stress triggering Dis-Arrest is removed (Fig. 1A - bottom panel), the broadly distributed recovery times result in the coexistence of growing and non-growing tolerant cells, leading to antibiotic persistence (Fig. 1B,C) [27, 25, 28]. As disrupted bacteria occur under many unrelated stresses, they may appear in many experimental conditions. Therefore, it is important to find ways to identify this subpopulation of cells at the single-cell level. In this study, we provide tools to analyze disrupted bacteria using single-cell and bulk RNA-seq, enabling a characterization of this ubiquitous phenotype. Furthermore, these tools should be generally relevant for the study of cells under stress.

**Figure 1:**
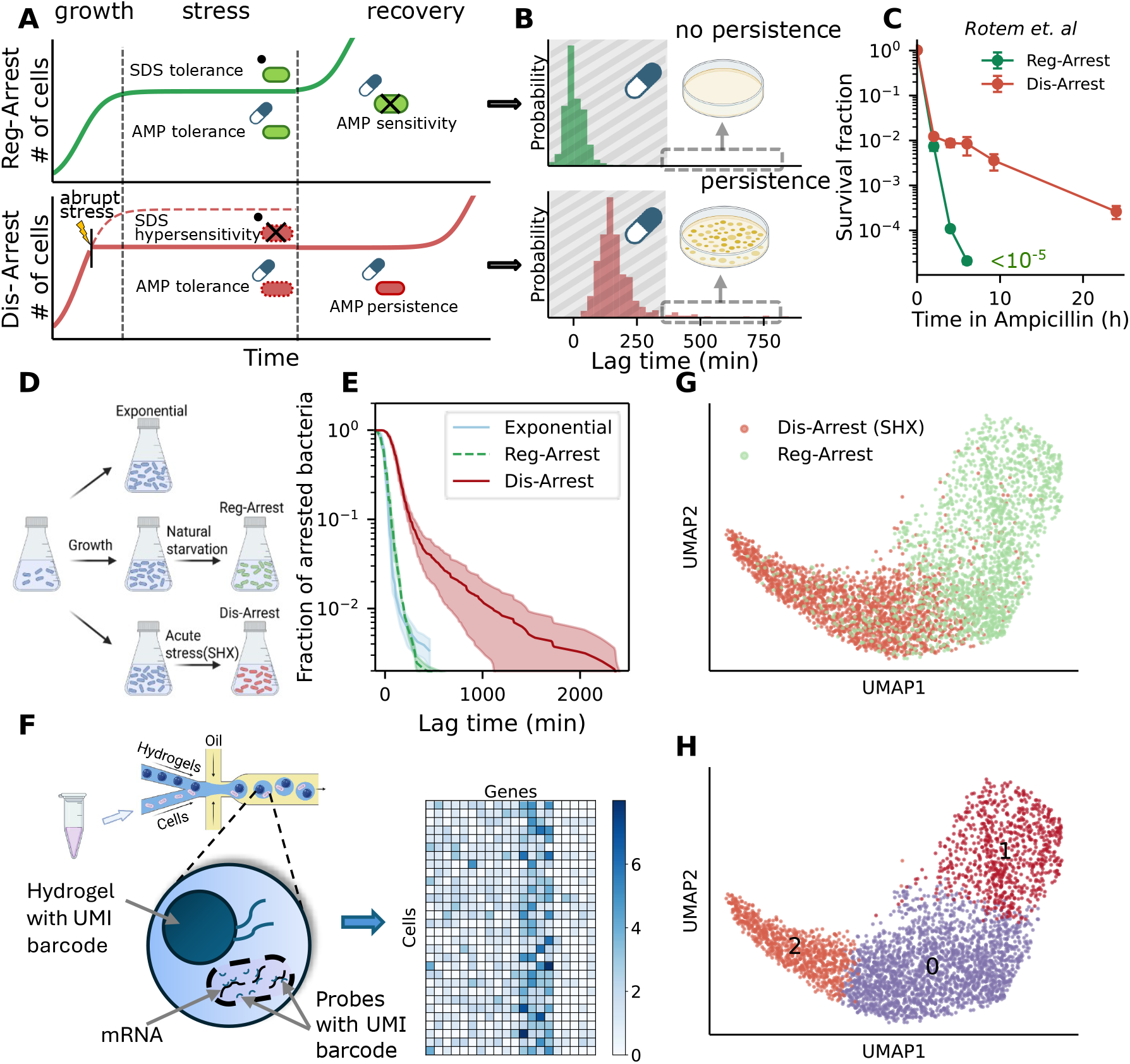
Characterizing the disrupted state (Dis-Arrest) using scRNA-seq. **(A)** Schematic growth curves of the bacteria highlighting the differences between natural starvation leading to Reg-Arrest and acute stress leading to Dis-Arrest during the 3 phases: growth, stress and recovery. **(B)** Empirical lag time distributions of cells diluted and regrown from exponential phase (top) or following acute stress (bottom), showing the typical long lag times and wide tail of cells recovering from Dis-Arrest (data obtained from Kaplan *et. al* [1]). The duration of antibiotic treatment is depicted by the shaded gray area. A sub-population of cells with lag times that exceed this duration can evade antibiotics and regrow, resulting in antibiotic persistence. **(C)** Kill curve of Reg-Arrest and Dis-Arrest bacteria treated with ampicillin during the recovery phase (reproduced from Rotem *et. al* [2]). Application of bactericidal antibiotics (ex. ampicillin) during the recovery phase produces persisters in the disrupted culture due to their long lag times. **(D)** Illustration of the experimental protocol. Exponential cells are harvested before applying stress. Reg-Arrest results from natural starvation. Dis-Arrest results from abrupt exposure to serine hydroxamate (SHX). Arrested samples are collected after 24 hours in growth-arrest. (**E**) Fraction of non-growing bacteria following regrowth of the bacterial cultures after Exponential, Reg-Arrest and Dis-Arrest conditions. The solid line and the ribbon represent the mean and confidence interval. Mean lag time and std over *n* = 3 biological replicates: Dis-Arrest 149±41; Reg-Arrest 1±13; Exponential −4±16. Statistical significance assessed by Welch’s t-test with a resulting Dis-Arrest to Reg-Arrest/Exponential p-value=0.02/0.01 respectively. **(F)** Schematic illustration of the single-cell RNA-seq experiment. A mixture of cells and probes is diluted and trapped in droplets containing unique cell barcodes. The pre-synthesized DNA probes target the mRNA transcripts, and contain unique molecular identifiers (UMI) and sequencing adapters. Finally, the resulting cell-gene count matrix is obtained after sequencing and mapping the reads to the *E. coli* genome. **(G)** UMAP plot of cells labeled by growth condition, representing the transcriptional profiles of the differently treated cells. **(H)** UMAP plot as in (G), with cells colored according to unsupervised clustering analysis of the gene expression data. Graphics created in biorender[118, 119].

The underlying mechanisms that control the tolerance and persistence following Dis-Arrest may differ greatly from those associated with Reg-Arrest. Several genetic factors have been shown to contribute to tolerance, typically attributed to a state of low translational activity related to regulated growth-arrest [29, 30, 31, 32, 33]. Although these findings provide valuable information, the lack of agreement between different studies on the genes involved led to the proposition that tolerance and persistence may result from the failure of various genetic programs rather than the activation of specific pathways [34, 1].

Previously, we have shown that prolonged lag times after stress removal, along with aging dynamics [1] and high sensitivity to environmental conditions [2], may indicate an underlying failure in the regulatory mechanisms of cells in Dis-Arrest. However, these physiological measurements do not provide direct information on the molecular noise, nor on the level of correlations between individual genes in Dis-Arrested cells. Gene-gene correlations can serve as a measurable proxy for regulatory interactions. For example, if gene A regulates gene B, their expression levels are dependent and they are expected to be either positively or negatively correlated. When this pathway is dysregulated, we should expect this correlation to be reduced [35]. If, indeed, Dis-Arrest originates from a loss of gene regulation rather than the activation of specific pathways, the signature of this dysregulation would be apparent in the loss of correlations between genes that belong to the impaired pathways.

In this study, our goal was to quantitatively determine the degree of dysregulation in gene expression in Dis-Arrest and to compare it to regulated cellular states. Non-adaptive stress responses have been shown to be associated with transcriptional dysregulation. Jensen et al. found that antibiotics decouple highly expressed genes from high-fitness genes across the metabolic network [36], and later it was shown, with a transcriptomic-entropy measure, that a more disordered global expression program predicts reduced fitness and increased antibiotic sensitivity across conditions [37]. Together, these results establish that the degree of organization of a stress response, rather than the activation of any single pathway, carries predictive information. However, the entropy measure requires a time-series of RNA-seq measurements over which gene expression changes, and is therefore not applicable to steady-states such as the regulated growth-arrest. To ask whether a culture is dysregulated from a single snapshot in time, we introduce GMP-Cor, a single-cell measure that quantifies the level of gene-gene coordination. Importantly, GMP-Cor can be applied to single time points, irrespective of whether the culture is at steady-state or not, and long after a perturbation. Achieving this requires single-cell resolution. Therefore, we perform single-cell RNA-seq on *E. coli* under different physiological conditions to compare transcriptional correlations in Dis-Arrest to those in Reg-Arrest.

Single-cell RNA sequencing (scRNA-seq) technologies offer a powerful method for analyzing gene expression diversity at the level of individual cells [38, 39, 40, 41, 42, 43, 44] and is required for evaluating transcriptional noise [45], or correlations between genes [35, 46, 47, 48, 49]. However, applying scRNA-seq to bacterial cells is challenging [50, 51, 52, 53, 54, 55]. The primary difficulty is the significantly lower levels of gene expression in bacteria, typically two orders of magnitude lower than in eukaryotic cells [56], requiring extremely sensitive experimental techniques to identify meaningful levels of transcripts. To effectively use scRNA-seq to characterize the cellular state by qualitatively evaluating gene-gene correlations, it is therefore essential to employ a high-throughput experimental technique that prioritizes per-cell sensitivity over maximizing the number of cells.

The reduced signal-to-noise ratio in bacterial single-cell studies also poses a major challenge for data analysis - the sparse data and high dimensionality of the transcriptomic space obscure true biological signals. Importantly, a proper mathematical approach that can mitigate these limitations is needed [57, 58, 45]. To this end, we develop a mathematical framework inspired by Random Matrix Theory (RMT) [59, 60, 61, 62] to model the gene-gene correlations. Instead of examining individual gene pairs, this approach offers a global measure of correlations [35]. Our analysis reveals that Dis-Arrest *E. coli* cells show significantly reduced gene correlations compared to cells in exponential growth or in regulated growth-arrest. Combined with insights from gene ontology (GO) analysis, these findings demonstrate that Dis-Arrest involves genome-wide dysregulation. Understanding the fundamentals of cells under stress should have broad implications for finding ways to either mitigate or enhance their global dysregulation. Finally, such control over dysregulation and its subsequent growth-arrest should help in devising strategies to fight persistent infections.

## 2 Results

### 2.1 Reduced Gene Expression Correlations in Disrupted Cells

To compare gene expression levels across different biological conditions, we perform scRNA-seq on *E. coli* cultures in three distinct states: exponential growth, regulated growth-arrest (Reg-Arrest), and disrupted growth-arrest (Dis-Arrest) (Fig. 1D). Exponentially growing cells are obtained by culturing *E. coli* for ∼ 13 generations to the exponential phase (OD < 0.1) to ensure balanced-growth and that there is no carryover from the previous stationary phase [15]. To induce growth-arrest, two distinct methods are used. For Reg-Arrest, cells are allowed to gradually deplete nutrients, leading to natural starvation and entry into the stationary phase. In contrast, Dis-Arrest is induced by abruptly exposing cells to serine hydroxamate (SHX), a chemical inhibitor that blocks charging of tRNA, causing unnatural serine starvation [21, 63, 64]. However, in a wild-type background, uncharged tRNA entering the ribosomal A site activates RelA and triggers (p)ppGpp synthesis, which broadly represses rRNA and ribosomal-protein transcription and results in a regulated arrest [11]. To reach a disrupted growth-arrest by SHX exposure, we use an *E. coli* strain which carries *relA1* and *spoT1* mutations leading to a relaxed stringent response. In this strain, abrupt SHX exposure arrests growth without allowing proper regulation, which is what makes the arrest disruptive. Importantly, adding SHX to cells already in Reg-Arrest reproduces the Reg-Arrest phenotype exactly: the lag-time distributions with and without SHX are indistinguishable (Extended Data Fig. S16A, adapted from Kaplan et al.[1]). This suggests that the important variable is the physiological state the cells are in when the stress arrives, and whether they have time to adapt.

As expected from Dis-Arrest cells [1], once the SHX is removed we observe a very broad and delayed distribution of lag times, in contrast to Reg-Arrest cells which rapidly and uniformly resume growth (Fig.1E). Moreover, as expected from the reductive growth characteristic of entry into regulated stationary phase, Reg-Arrest cells are ∼2x smaller than exponentially growing cells. In contrast, Dis-Arrest cell sizes are similar to exponentially growing cells (Extended Data Fig. S13A,C). Our interest is to characterize these growth-arrested states when Dis-Arrest is clearly different than Reg-Arrest by physiological measurements, namely after about 24 h at growth-arrest (Methods). Because the synchrony of the arrest matters for interpreting the population as a single state, we measured it directly by time-lapse imaging in a mother machine, scoring every elongation and division event (Extended Data Fig. S14, Methods). We observed a uniform division arrest with no division events occurring more than 40 min after SHX addition.

Cultures at either Reg-Arrest or Dis-Arrest were analyzed with scRNA-seq using ProBac-seq [55]. Probac-seq utilizes libraries of pre-synthesized DNA probes, thus removing the rRNA depletion step necessary in most bacterial single cell techniques available (see Fig. 1F). This is beneficial when studying starved bacteria, which have even lower mRNA counts than growing bacteria [2], as the signal could otherwise be obscured by residual rRNA sequencing. The scRNA-seq results are then analyzed to understand gene expression profiles and correlation patterns across different growth conditions.

To visualize the transcription profiles of the single cells we use a uniform manifold approximation and projection (UMAP) [65] embedding of the combined dataset from the two growth arrested conditions - Reg-Arrest and Dis-Arrest (Fig. 1G-H). Figure 1G shows the mapping of the different growth conditions to the 2-dimensional manifold, while figure 1H assigns colors to the same UMAP plot according to an independent unsupervised clustering of the single cells. Analysis of the marker genes of each cluster reveals a few prominent features of the growth-arrested states (Extended Data Fig. S7, for a full list of marker genes see Supp. table 1). Cluster 1, which is predominantly occupied by Reg-Arrest cells, is characterized by a reduction in many ribosomal and translation related genes (*rplT, rplE, rpsB, rpsC, …*), indicating an adaptive downregulation of translation aimed at conserving resources under stress [63, 66]. The cluster of cells exposed to SHX (cluster 2) stood out by exhibiting robust up-regulation of the phage shock protein (*psp*) operon. While the *psp* operon is typically strongly induced by envelope-damaging perturbations (heat, osmotic shock, filamentous phage, alkaline stress) [67, 68], it is not canonically associated with the response to nutrient starvation [64]. We hypothesize that its upregulation may be an indirect effect of dysregulation. This is consistent with recent observations, showing that different conditions leading to Dis-Arrest result in the loss of membrane homeostasis and increased membrane permeability [2].

In order to evaluate whether Dis-Arrest bacteria are indeed dysregulated, we now turn our attention to the network of gene interactions. Our goal is to compare the level of global correlations in gene expression across the three conditions mentioned above. When assessing the gene-gene correlation levels in the single-cell data, it is important to recognize that any straightforward calculation of the pairwise correlation between genes is affected by “spurious correlations”. Even random uncorrelated data of high dimensionality with a limited number of samples would display such spurious correlations, as shown in Fig. 2A. Therefore quantifying genuine correlations (those that arise due to true dependencies) in experimental data necessitates a method capable of effectively excluding the spurious correlations from the analysis. To address this, we first derive an analytical model, based on the eigenvalue distribution of the correlation matrix [69, 59]. If the correlations in normally distributed random data are analyzed, the resulting eigenvalue distribution is the well known Marchenko-Pastur (MP) distribution [70] (Fig. 2A). Our model generalizes the MP distribution to describe the eigenvalue distribution of data with genuine correlations - hereinafter termed the *Generalized MP* (GMP) distribution. This distribution is tuned by a single parameter χ (Extended Data Fig. S1, Methods and Supp. note 1 for a detailed derivation). Figure 2**B** displays the original MP distribution (χ=0) and the GMP distribution for various values of χ. When comparing the distribution of random data (MP) to the distribution of correlated data (GMP), the presence of correlations becomes apparent in two distinct regions (Fig. 2B). The first is an extended tail in the probability distribution, reflecting a greater likelihood of large eigenvalues beyond the range predicted by the standard MP distribution. A second feature of the GMP distribution is the increased probability of small eigenvalues. These two features in the eigenvalue distributions provide insight on how genuine correlations are expressed beyond the impact of spurious correlations caused by finite-size effects (Fig. 2B).

**Figure 2:**
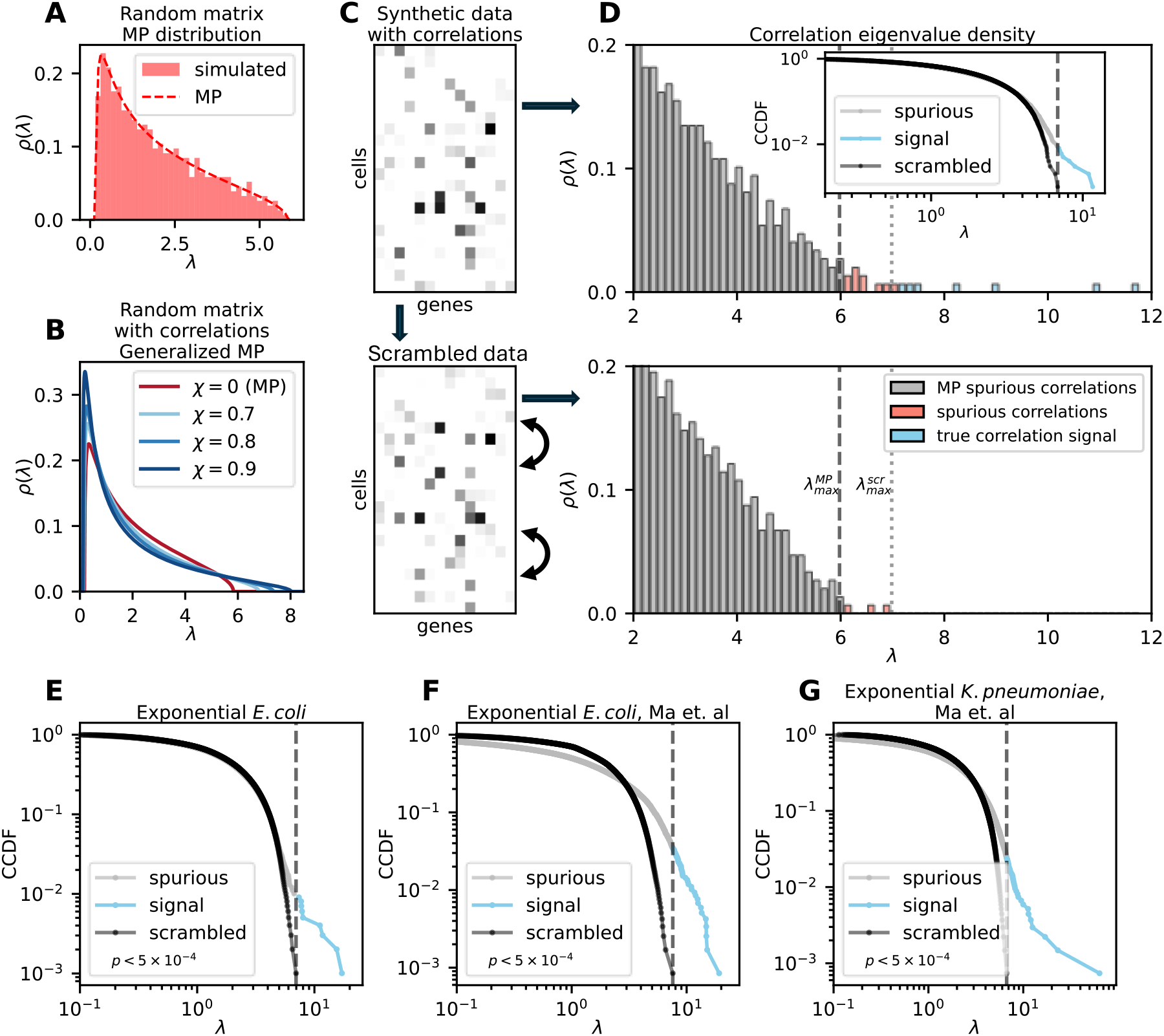
Identifying correlations in the *E. coli* transcriptional network from the eigenvalue distribution of the single-cell correlation matrix. **(A)** Correlation eigenvalue spectrum of a random Gaussian matrix with a dimensionality to sample size ratio *α* = *P/N* = 2. The eigenvalue density of this matrix is shown along with the analytical prediction of the Marchenko-Pastur (MP) distribution. **(B)** Analytical plot of the Generalized Marchenko-Pastur (GMP) distribution for *α* = 2 and different values of the correlation strength parameter *χ*, illustrating how increasing *χ* progressively shifts the weight of the distribution toward larger eigenvalues. **(C)** Representative synthetic sparse count matrix (top), generated with strong gene-gene correlations and its scrambled counterpart (bottom). Scrambling along the columns preserves gene distributions and overall data structure while erasing gene-gene correlations. **(D)** Eigenvalue density of the synthetic matrix (top) color-coded by region: grey bars below the MP upper edge 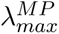(dashed line) indicate dimensionality induced noise. Red bars between 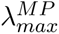and the scrambled maximum 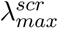(dashed line) reflect spurious correlations. Blue bars exceeding 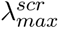represent meaningful correlations. Inset: complementary cumulative distribution function (CCDF) o log-log axes of both original and scrambled eigenvalue distributions. The threshold 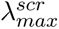(dashed line) separates spurious (grey) from true correlation signal (blue). The scrambled eigenvalue density (bottom) provides the empirical noise ceiling. **(E-G)** Log-log CCDF of eigenvalues for exponential *E. coli* from this study **(E)**, exponential *E. coli* **(F)** and *K. pneumoniae* **(G)** from Ma *et. al* [54]. In all CCDF panels grey represents spurious eigenvalues, blue are correlation signal eigenvalues above 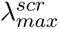and black is the scrambled distribution. Statistical significance of the original vs. scrambled distributions is obtained by a permutation test (methods) with the resulting p-value annotated in each plot.

While the analytical GMP distribution provides crucial insight on how to analyze the distribution of correlation eigenvalues, it is far from being representative of scRNA-seq data. Real data is notoriously sparse, and individual genes span widely in both their mean expression and cell-cell variability. To address this, we generate synthetic scRNA-seq data with underlying correlations (Fig. 2C, Extended Data Fig. S2, Supp. Note 2). This numerical model allows us to tune the level of underlying correlations by a single parameter χ and observe the resulting correlation eigenvalue distributions in data that mimics the nature of real experimental data. The eigenvalue distribution is shown in Fig.2D, along with the distributions computed from surrogate scrambled data. This scrambling of the data represents full dysregulation and therefore removes correlations, providing a clear comparison of the original data to a null distribution of the same structure but without correlations. Therefore, we interpret the eigenvalues above the maximal scrambled eigenvalue as the true correlation signal. The inset shows the same distributions as a log-log complementary cumulative distribution function (CCDF) to better visualize the tail of the distributions (Fig. 2D). The results of this simulation confirm that the extended tail of large eigenvalues is caused by genuine correlations.

We now turn to the analysis of gene-gene correlations in the scRNA-seq datasets, starting with the analysis of exponentially growing cells. Exponential growth requires the coordinated action of many genes and is therefore highly regulated, which is expected to result in a high level of gene correlations. Figures 2E-G show the eigenvalue distributions of the correlation matrices of scRNA-seq calculated from our data, as well as from datasets in two additional studies of exponentially growing, untreated bacterial cultures (Fig. 2E: data from this study, Fig. 2F: *E. coli* data from Ma et. al [54], and Fig. 2G: exponential *K. pneumoniae* cells from Ma et. al [54]). Despite differences in experimental protocols, bacterial strains, and single-cell technologies, all datasets consistently exhibit deviations from their scrambled counterparts. This shift reflects the expected gene-gene correlations present in unperturbed, exponentially growing bacteria.

Turning to bacteria under stressful conditions, we apply the same methodology to Reg-Arrest and Dis-Arrest cells. Figures 3A and 3B display the eigenvalue distributions of the correlation matrices for Reg-Arrest and Dis-Arrest cells, respectively, together with the corresponding results from surrogate scrambled data. For comparison, Fig. 3C,D show the simulated data results for strong and weak correlations respectively. To quantify the strength of true correlations in the data, we define the GMP correlation index (GMP-Cor) as the sum of eigenvalues above the maximum scrambled eigenvalue (see Methods). The large eigenvalues represent genes that are correlated between cells, therefore the GMP-Cor metric enables us to quantify the overall degree of gene-gene correlations in each dataset. This analysis shows that Dis-Arrest datasets exhibit weaker correlations compared to either Reg-Arrest or Exponential growth datasets. We summarize the GMP-Cor values for all datasets analyzed in this study showing that this metric provides a statistically significant classification between disrupted and regulated conditions (Fig. 3E, Extended Data Fig. S3, Supp. Note 4). We extend the use of the simulated data generator presented above to calibrate the GMP-Cor readout against data with a known ground truth correlation strength. Sweeping χ from 0 to 1 at fixed sparsity, yields a calibration curve (Extended Data Fig. S4A). This defines the expected ranges for dysregulation (low), partially regulated (intermediate) and regulated (high) states. The calibration curve gives us important insight: we observe that GMP-Cor is a robust evaluation of the correlation strength, giving similar results for synthetic data that mimics the real data structure, as well as for the theoretical random matrix analysis of the GMP distribution (dashed curve in Extended Data Fig. S4A).

**Figure 3:**
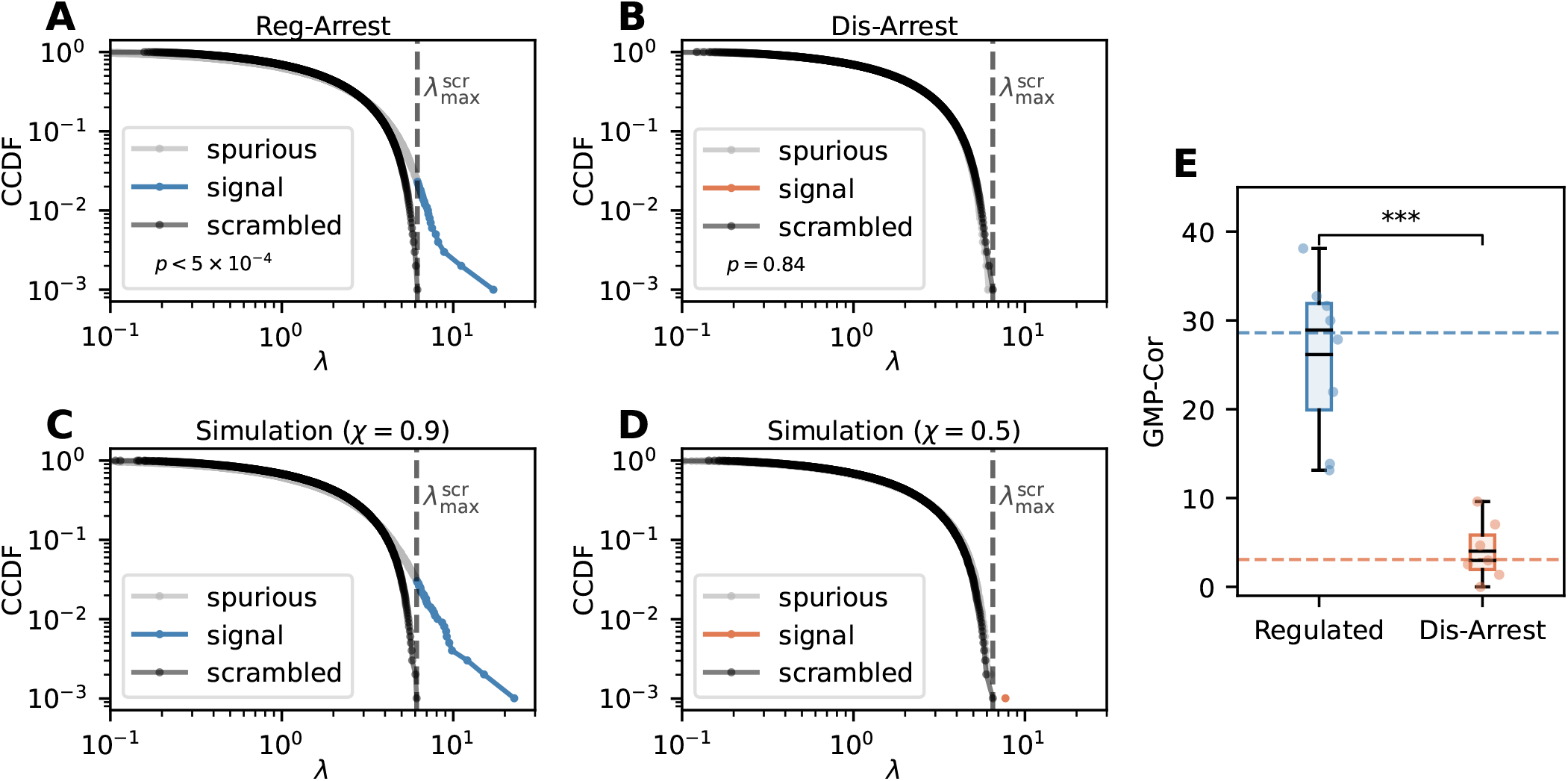
The disrupted state (Dis-Arrest) exhibits significantly weaker transcriptomic correlations than regulated (Reg-Arrest) cells. **(A-B)** Log-log CCDF of correlation eigenvalues for Reg-Arrest **(A)** and Dis-Arrest **(B)** experimental conditions. The dashed vertical line marks 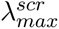- eigenvalues exceeding this threshold (colored) reflect biologically meaningful gene-gene correlations absent in the scrambled control (black). Statistical significance of the original vs. scrambled distributions is obtained by a permutation test (methods) with the resulting p-value annotated in each plot. **(C-D)** Corresponding CCDFs from synthetic data simulations at correlation strength *χ* = 0.9 **(C)** and *χ* = 0.5 **(D)**, qualitatively reproducing the high and low correlation profiles observed experimentally. **(E)** Distribution of GMP-Cor values across individual samples in the Regulated and Dis-Arrest groups. Individual data points are overlaid on the box plots. Horizontal dashed lines represent the medians of each group. Statistical comparison by two-sided Mann-Whitney U test (\*\*\**p <* 0.001).

Analyzing the correlation eigenvectors shows that the highly correlated modes are broadly spread across the transcriptome and contain interpretable biology. The leading modes are dominantly enriched for translation related pathways in all conditions, motility and biosynthesis in exponential cells, and lipid biosynthesis and stress response in both Reg-Arrest and Dis-Arrest. This analysis suggests that the genes that carry the regulated/dysregulated signal are similar, but the level of regulation is expressed in the magnitude of the eigenvalues, which is exactly the readout of GMP-Cor (see supp. note 3). Importantly, the correlation spectrum readout behaves differently from bulk measures of variability when a population is heterogeneous: a mixture of two subpopulations is averaged by bulk data, resulting in a loss of structure, but raises the GMP-Cor signal in single cell data, because the mixture itself generates a coordinated mode separating the two groups (Extended Data Fig. S6A-B). A reduced correlation signal therefore cannot be produced by subpopulation structure, and specifically indicates a loss of coordination within cells. Taken together, our results reveal a loss of gene-gene correlation and global dysregulation of the Dis-Arrest bacteria.

### 2.2 Evidence of Dysregulation in Disrupted Cells from Gene Ontology Analysis

The GMP distribution provides a universal framework that assumes a simplified structure of the gene network. However, true gene interactions display non-universal characteristics linked to specific gene clusters responsible for distinct biological functions. Gene Ontology (GO) analysis identifies these functional gene groups and utilizes them to detect biological behavior differences between cells under different conditions. GO analysis regroups the accumulation of knowledge on many datasets and represents the gene expression programs expected from cells in regulated states, whether growing or non-growing. Therefore, dysregulation should also be apparent in deviations from the expected enrichment of GO terms, according to the specific physiological condition.

To test whether GO analysis can confirm the dysregulation observed in the scRNAseq data in Dis-Arrest, we analyze bulk RNAseq in the same conditions: Reg-Arrest and Dis-arrest compared to exponentially growing cells before growth-arrest (see Methods). Due to the higher sensitivity of the bulk measurements, more subtle differences in expression patterns can be identified. Differentially expressed genes (DEG) were assessed from the data by comparing either Reg-Arrest or Dis-Arrest to exponential cells (Methods). DEG analysis was consistent with the scRNAseq data, identifying a strong upregulation of the *psp* operon in Dis-Arrest, as well as a reduction of ribosomal genes in the Reg-Arrest condition. Further, per-gene mean expression agrees between the two data types (Dis-Arrest scRNA-seq vs bulk: Pearson r = 0.75, Exponential: Pearson r = 0.69, Reg-Arrest: Pearson r = 0.34; Extended Data Fig.S11). Perfect agreement is not expected, since the single-cell protocol relies on a pre-synthesized probe set whose composition and capture efficiency differ from bulk RNA-seq. The DEG sets from Reg-Arrest and Dis-Arrest cells were then analyzed for enriched GO terms to assess whether specific biological functions are activated (see Methods, for a full list of enriched GO terms see Supp. table 3). By representing each GO term as a subset of all genes in the genome, the degree of overlap between DEGs and GO terms serves as a measure of the likelihood that a particular genetic pathway is involved. In a properly functioning cell, DEGs are expected to significantly overlap with relevant GO terms. In contrast, in a fully dysregulated state, the set of DEGs would contain random elements from the large genome space, resulting in minimal overlap with any specific GO term. This overlap can be reduced further by a loss of coherence, when genes from the same pathways are differentially expressed in opposite directions. An example that illustrates this concept on the chemotaxis GO term is shown in Fig. 4A. The level of coherence can be understood here as the relative amount of significantly down-regulated genes in this group. While the Reg-Arrest column is strongly coherent (all blue), the Dis-Arrest column displays both up (red) and down regulated (blue) genes - indicating that this group of genes is not coherently turned off. We show the chemotaxis GO term here because it clearly highlights the coherence effect - the same qualitative contrast holds for any term with a pronounced difference in enrichment score which we present below. The degree of overlap and coherence can then be quantified for each GO term with a p-value obtained from a GO enrichment test. These p-values quantify the deviation of the current cellular state from maximal dysregulation, analogous to the null hypothesis of no functional association. Below we use this approach on all the significantly downregulated GO terms to evaluate the level of coherence in gene expression in bulk transcriptomics between Reg-Arrest and Dis-Arrest.

**Figure 4:**
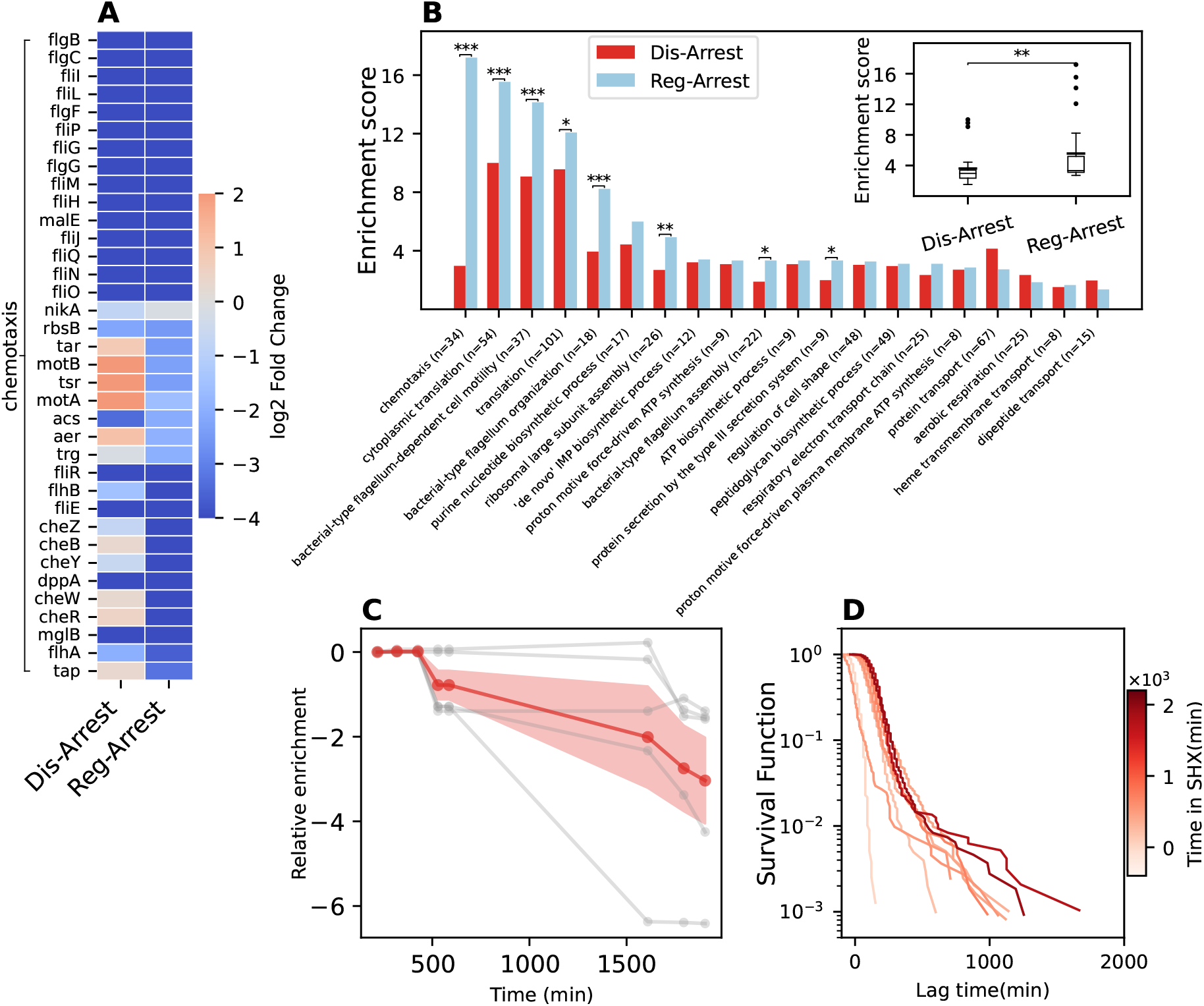
GO analysis shows stronger coherence of terms in the regulated state (Reg-Arrest) compared to the disrupted state (Dis-Arrest). **(A)** Heatmap of the log_2_ fold change of gene expression in Dis-Arrest and Reg-Arrest compared to exponential cells. To illustrate the concept of coherence, we show genes associated with the chemotaxis GO term. While the Reg-Arrest column contains only downregulated genes (shades of blue), the Dis-Arrest column exhibits also upregulated genes (shades of red). This indicates that chemotaxis is more coherently downregulated in Reg-Arrest than in Dis-Arrest. **(B)** Enrichment score (− log_10_(*p_adj_*)) of common enriched GO terms. Red bars represent Dis-Arrest, and blue bars represent Reg-Arrest. Higher bars represent a larger enrichment score, and consequently, stronger coherence. Statistical significance of the difference between conditions for each GO term is based on a random resampling test. \**p <* 0.05; \*\**p <* 0.01; \*\*\**p <* 0.001. Inset: Box plot comparing the distributions of enrichment scores for the same number of most significant GO terms that are enriched in each study. The box represents the first to third quartiles. The solid black line denotes the mean, and the circular markers are outliers. The Mann-Whitney U-test confirms that the 2 distributions are significantly different with a resulting p-value of 3.1 × 10*^−^*^3^. **(C)** Relative enrichment 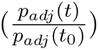of significantly enriched GO terms at different time points during SHX treatment, showing an overall trend of reduced GO term coherence over time (the y-axis is in − log_10_ scale so negative values represent reduced enrichment). DEG’s are compared to the expression levels at the moment of SHX application. The red markers and ribbon represent the mean and ±*σ* confidence interval of the relative enrichment values across all significant GO terms. **(D)** Survival function of lag times after recovery from different treatment durations in SHX, measured on the same culture as in (C). The broad distributions along with the strong dependence on treatment duration are canonical characteristics of the aging dynamics in Dis-Arrest[1].

Both Reg-Arrest and Dis-Arrest experience similar environmental stress (starvation), and therefore unsurprisingly do share several commonly enriched GO terms among their down-regulated genes (i.e. chemotaxis, translation, flagellum organization). However, the key difference between these states lies in the degree of certainty with which these biological functions are expressed, namely the level of coherence in gene expression modules, as quantified by the enrichment score (negative log_10_ of the adjusted p-value (p*_adj_*) obtained by the GO enrichment test). We note that all GO terms that have a significantly different enrichment score between the conditions, are more enriched in Reg-Arrest compared to Dis-Arrest (Fig. 4B). Furthermore, when evaluating all significant GO terms specific to each condition, Dis-Arrest consistently exhibits lower enrichment scores, indicating weaker functional regulation across both shared and unique GO terms (Fig. 4B inset). To relate our conditions to the established framework of the core stress response, we asked whether the RpoS regulon behaves differently in Reg-Arrest and in Dis-Arrest. We annotated RpoS-regulated genes (RegulonDB) in the same DEG data and tested their enrichment (Extended Data Fig. S9A-B). Reg-Arrest shows a pronounced up-regulation of the regulon, as expected for a controlled entry into stationary phase. SHX-induced Dis-Arrest shows only a weak, marginally significant up-regulation, consistent with the lower coherence observed in GO terms.

To examine how dysregulation develops over time under Dis-Arrest conditions, we performed bulk RNA-seq experiments on samples collected at multiple time points during SHX treatment (methods). GO analysis of these datasets reveals a reduction of coherence over time for all consistently enriched GO terms (Fig.4C). This indicates that over the course of stress exposure, cells lose biological organization, consistent with the increase in cellular disruption over time inferred by measuring the lag time distributions on the same culture (Fig. 4D) [1, 71, 72]. In contrast, analysis of an intermediate time-point in Reg-Arrest (480 min) reveals that there is no overall reduction of GO term coherence in Reg-Arrest, but rather most GO terms increase their enrichment score over time (Extended Data Fig. S10). Thus, we can link the Dis-Arrest observed in Kaplan et al. [1] and its strong dependence on stress duration, to the dynamics of dysregulation during stress exposure. This analysis of coherence in GO terms provides further support for the identification of global dysregulation in Dis-Arrest, as demonstrated in the single-cell data.

### 2.3 Loss of Gene Correlations in Growth-Arrested Cells Induced by Toxin Exposure

To further investigate the role of dysregulation in tolerance, we expanded our analysis to a well-characterized stress condition known to induce high levels of antibiotic tolerance and persistence, namely the induction of a toxin-antitoxin module [29]. For this purpose, we engineered a bacterial strain with a chromosomally integrated inducible toxin gene, *vapC* [73]. The toxin VapC is a ribonuclease that cleaves the fmet-tRNA, inhibiting protein synthesis, consequently causing growth-arrest and increased persistence [74, 75, 76, 77] (typical lag time distributions are shown in Fig. 5I). Interestingly, the VapBC toxin-antitoxin module was involved in mutations frequently emerging during evolutionary adaptation under cyclic antibiotic exposure, and resulting in a very high level of antibiotic tolerance[78, 79]. In addition to its direct relevance in antibiotic response, we chose VapC for two main reasons. First, VapC-induced arrest had not previously been characterized as Reg-Arrest or Dis-Arrest, making it an opportunity to test the metric. Second, the VapC toxin arrests growth through a completely distinct mechanism from SHX, allowing us to test our hypothesis beyond the framework of the stringent response. Our goal was to determine whether the growth-arrest induced by VapC expression results in Dis-Arrest or Reg-Arrest, using the GMP-Cor metric for quantifying dysregulation.

**Figure 5:**
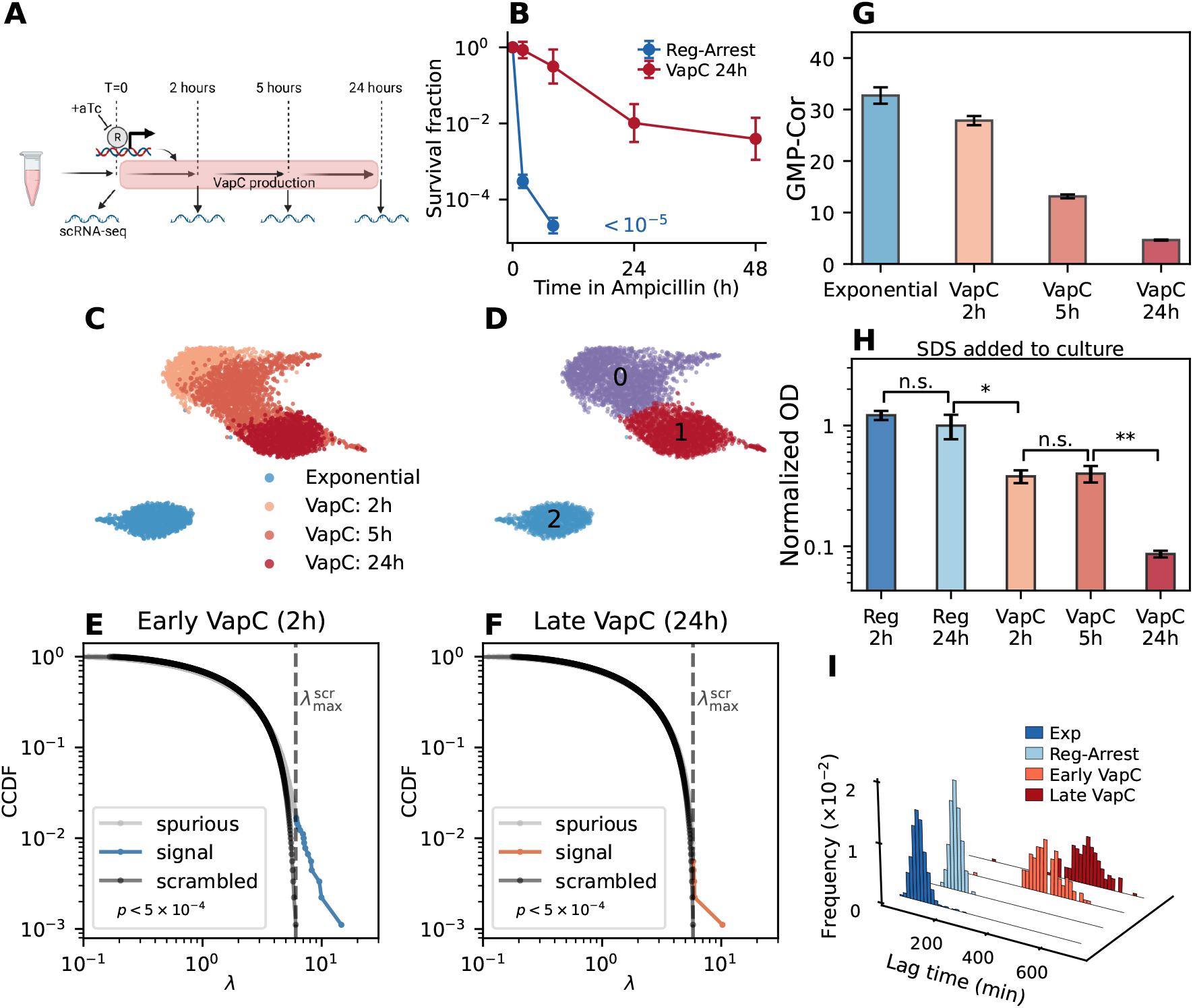
Long time exposure to VapC reduces transcriptomic correlations. **(A)** Schematic diagram of the experimental procedure. Cells with an inducible promoter upstream of the *vapC* gene are grown to mid-exponential phase. At *t* = 0 aTc is added to the medium promoting VapC production. After 2, 5 and 24 hours, samples are taken from the culture for scRNA-seq measurements. **(B)** Kill curve of VapC induced cells following 24h of VapC induction or Reg-Arrest upon dilution into fresh medium with Ampicillin. Error bars represent the geometric SD of *n* = 3 biological replicates. **(C)** UMAP plot of cells labeled by treatment duration. **(D)** Cells are labeled on the UMAP plot presented in (C) by the results of unsupervised clustering analysis. **(E-F)** Log-log CCDF of correlation eigenvalues for VapC induced cells after 2 hours **(E)** and 24 hours **(F)** of VapC induction. The dashed vertical line marks 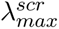- eigenvalues exceeding this threshold (colored) reflect biologically meaningful gene-gene correlations absent in the scrambled control (black). Statistical significance of the original vs. scrambled distributions is obtained by a permutation test (methods) with the resulting p-value annotated in each plot. **(G)** GMP-Cor values of the datasets without VapC induction (exponential) and VapC induction for 2, 5 and 24 hours, representing the difference in correlation strength between early and late times of VapC induced stress. Error bars are ±SD obtained from a permutation test (methods). **(H)** OD measurements of the different cultures incubated with SDS 1% for 20 hours. Values are normalized by OD just before treatment. The bars show that only VapC cultures are affected by the treatment (an OD decrease is understood as cell lysis) and that long-time VapC cultures are significantly more sensitive than both early VapC and Reg-Arrest. Statistical significance is assessed by a one-sided t-test performed on *n* = 3 biological replicates for VapC cultures and *n* = 2 biological replicates for Reg-Arrest cultures. Two technical replicates are measured for each biological replicate. **(I)** Normalized histograms of the regrowth statistics (lag times) for exponential (Exp) Reg-Arrest and VapC production for short (*t* = 240min) and long (*t* = 1400min) times. Exp and Reg-Arrest are plotted in blue, and cells that undergo VapC induction are plotted in red. Graphics created in biorender[120].

We induced the production of VapC during the exponential growth phase, resulting in growth-arrest (Methods). To establish directly that this condition produces persisters, we measured survival rates under ampicillin during the recovery phase (Fig. 5B). We observe biphasic killing and significantly enhanced survival after 24h of induction relative to the regulated control. We performed scRNA-seq on the growth-arrested *E. coli* cells at 2, 5 and 24 hours after VapC induction (Fig. 5A). A distinct transcription profile in the UMAP embedding is apparent for the exponentially growing cells (before VapC induction), followed by a continuous transition between 2 and 24 hours of VapC production (Fig. 5C). Unsupervised clustering on the gene expression data identifies three distinct cell states (Fig. 5D). Exponential cells are found in cluster 2, which exhibits expression of flagellar genes, protein synthesis and metabolic genes - hallmarks of a motile, rapidly growing phenotype. Clusters 0 and 1 capture early and late responses to VapC, respectively. Cluster 0, dominated by early post-induction cells, shows elevated expression of ribosomal genes, suggesting sustained protein synthesis. In contrast, Cluster 1, enriched with late (24 hours) post-induction cells, exhibits a metabolic shift toward nutrient scavenging and biosynthesis, along with markers of DNA repair and oxidative, heat, and envelope stress responses (Extended Data Fig. S8, for a complete list of cluster-specific marker genes, see Supp. table 2). Notably, cells collected 5 hours after induction are distributed between clusters 0 and 1, indicating a gradual phenotypic transition around this time point.

To determine whether each of these phenotypes can be associated with Dis-Arrest, we search for signatures of dysregulation in their gene expression profiles. The correlation spectrum of VapC-induced cells reveals a signature of gene-gene correlations within 2 hours after induction (Fig. 5E), followed by a pronounced loss of correlations after 24 hours (Fig. 5F). This trend is quantified by the corresponding GMP-Cor values (Fig. 5G). The lower GMP-Cor value 24 hours after VapC induction suggests that prolonged VapC induction pushes cells into Dis-Arrest, similarly to the growth-arrest induced by SHX.

The scRNA-seq data suggests that VapC induction initially triggers a regulated response during the first few hours, which later degrades into a disrupted growth arrest. The long lag times during the early VapC time points (Fig. 5I) are not a contradiction to this conclusion, because VapC accumulates in the cells and creates a long delay even after a short induction. The lag is therefore bounded from below by the time required for the toxin to be cleared, independent of the regulatory state.

To test the establishment of the disrupted arrest after 24 hours of toxin induction, we performed physiological measurements on the VapC expressing cultures. Specifically, we assessed cellular membrane integrity, as membrane impairment is a characteristic feature of disrupted growth arrests induced by various conditions[2]. For this purpose, we test the integrity of the cell’s membrane after VapC induction. The low GMP-Cor value for cells under prolonged VapC stress suggested that these cells would be at Dis-Arrest and therefore be susceptible to treatments that would otherwise not penetrate the *E. coli* membrane. One such substance is sodium dodecyl sulfate (SDS), an anionic surfactant that is mainly used to lyse cells [80]. It was shown that whereas Reg-Arrest and exponentially growing *E. coli* cells are essentially unaffected by 1% SDS [81, 82], the same treatment on Dis-Arrest cells induced by SHX results in lysis [2]. Consistent with the detection of dysregulation in cells after 24h of VapC induction, we also observe a markedly increased sensitivity to 1% SDS in these cells, whereas control Reg-Arrest cells were largely unaffected (Fig. 5H). Furthermore, single-cell imaging shows that VapC arrested cells are approximately twice the size of their Reg-Arrest controls, similarly to SHX-induced Dis-Arrest (Extended Data Fig. S13A).

Together with the predictions of the GMP-Cor metric, the SDS assay and cell-size analysis provide independent evidence that extended VapC induction drives cells into a Dis-Arrest state. These results demonstrate the power of the GMP-Cor metric to identify the global dysregulation in Dis-Arrest, not only when triggered by SHX, but also following a very different stress inflicted by VapC expression.

## 3 Discussion

In this study, we characterized the disrupted (Dis-arrest) *E. coli* cells by quantifying the level of dysregulation using both bulk and single-cell RNA-seq. Antibiotic tolerance observed in regulated growth-arrested bacteria arises from entry into growth-arrest mediated by dedicated gene regulatory networks. In contrast to this behavior, we show that the antibiotic tolerance associated with Dis-Arrest is governed by dysregulation. We quantify this dysregulation by introducing GMP-Cor, a measure of the overall gene-gene correlation strength. A quantitative approach to study non-adaptive, disordered stress responses has previously been proposed through transcriptomic entropy [37]. However, GMP-Cor addresses a different question on different data. Transcriptomic entropy is a bulk readout that characterizes how broadly cells sample expression space shortly after a perturbation, and it requires a defined reference distribution and time series. GMP-Cor, by contrast, is a single-cell, steady state measure of whether the gene–gene correlation structure is preserved relative to a scrambled null, long after a perturbation, and can identify dysregulation also at the sub-population level. On the other hand, GMP-Cor requires single-cell data, which remains less accessible than bulk measurements, particularly in bacterial systems.

We demonstrate a loss of gene–gene correlations under two distinct conditions that lead to Dis-Arrest, providing direct evidence that these antibiotic-tolerant cells exhibit dysregulation of their gene regulatory network. We note that low values of GMP-Cor do not necessarily mean zero regulation, as they can translate to a moderate correlation strength (see Extended Data Fig. S4A). However, the detected significant decrease of GMP-Cor in Dis-Arrest compared to regulated states supports the view that Dis-Arrest is characterized by chaotic-like dynamics [83], that account for the observed heterogeneity and broad distribution of regrowth lag times after stress removal [1].

The distinction between Dis-Arrest and Reg-Arrest can be understood by analogy to the differences between necrotic and senescent states in mammalian cells. Both states are typically accompanied with cell-cycle arrest. However, whereas senescence is an organized regulatory program aiming at maintaining mammalian cells at an irreversible cell-cycle arrest without death [84], necrosis is often the result of acute stress leading to uncontrolled growth-arrest, and loss of homeostasis followed by death [85]. Dis-Arrest bacteria have features similar to necrotic cells, such as dysregulation and damaged membranes [86]. However, unlike necrotic mammalian cells, which are doomed to die, Dis-Arrest bacteria exhibit a remarkable resilience that enables them to eventually re-enter the cell-cycle once the stress is removed.

Our new metric for dysregulation (GMP-Cor) provides a global measure of correlations on the scRNA-seq data and can be used as a general “quality control” test for how regulation dominates the transcriptional response. This analytical tool, complemented by numerics, addresses the challenges of low signal-to-noise ratios in scRNA-seq and the spurious correlations inevitably arising in typical data sets. Although it allows to identify correlation patterns in sparse data, this tool does not come without limitations. Our analysis was performed on datasets with a typical mean of ∼ 50 − 200 detected genes per cell. It is important to note that applying this technique to datasets with substantially lower gene detection rates may not provide meaningful results. Clearly, in the limit of very few genes per cells, the ability to detect gene-gene correlations goes down. Further, GMP-Cor is influenced by sample coverage or depth, necessitating a careful filtering of cells to compare conditions on an even ground (Methods). Nevertheless, we do not observe a significant correlation between the gene detection rate and GMP-Cor within the same experimental technique (Extended Data Fig. S5), meaning that our measure of correlations is not determined from sequencing depth alone. While our approach is powerful for the comparison of global correlations between physiological conditions, we note that it may have limitations for comparing samples obtained from different experimental techniques. This is due to the fact that different sources of technical variability (sequencing depth, cell filtering, etc.) have a major effect on the signature of correlations in the data. Here, we were careful to compare Dis-Arrest and regulated cells within the same experiment.

Additional support for the power of the GMP-Cor metric in detecting dysregulation was found in the loss of gene correlations determined by enrichment of GO terms. We analyzed bulk RNA-seq in the same conditions and observed a loss of coherence of enriched GO terms in Dis-Arrest compared to Reg-Arrest cells. This loss of coherence, which is manifested in lower enrichment scores, can be interpreted as a deviation from a controlled activation of stress response - an expected effect in a state of genome-wide dysregulation. Although this approach offers a rather indirect measure of dysregulation in comparison to our single-cell analysis, it has the advantage of relying on bulk measurements, which are more accessible and tend to produce higher quality data. While the GMP-Cor metric of the single-cell data provides information on the global level of correlations in the gene network, it potentially overlooks small gene clusters that are expected to be highly correlated and are represented by the GO terms. By focusing on these clusters, the analysis of GO term coherence complements our single-cell analysis, providing a broader picture of dysregulation.

The Dis-Arrest conditions studied here are fundamentally different in the way they disrupt cellular growth (SHX-induced starvation and VapC-induced tRNA cleavage), and differ in the pathways they activate. SHX-induced starvation leads to an enhanced expression of the *psp* module, while VapC induces metabolic shifts towards alternative carbon sources and increased biosynthesis[73]. Furthermore, SHX-induced Dis-Arrest upregulates RpoS controlled genes, and VapC-induced arrest - which has no a priori relation to the stringent response - does not activate this program (Extended Data Fig. S9C). As expected, they do share a core stress-response. First, a general protective response to protein aggregation (e.g. upregulation of the AAA+ chaperones *clpA*/*clpB* and the *hslO* /*hslV* genes) [87, 88]. Second, both treatments induce factors that aim to maintain envelope integrity - notably the major lipoproteins Lpp and LolB and the TolC efflux channel - suggesting a common challenge to maintain membrane homeostasis [89, 90]. In other words, Dis-Arrest is not a single phenotype, not even a defined “state”, but rather a cellular condition that exhibits dysregulation across different treatments. These observations are supported by the reduced membrane integrity in both SHX-induced Dis-Arrest [2] and VapC induced growth-arrest demonstrated by the SDS sensitivity test in Fig. 5H.

The dysregulation that we observe in Dis-Arrest cells can be understood in two conceptually different frameworks for the dynamics of cells under acute stress. In the first, evolution plays a major role in designing a mechanism that enhances variability. This is achieved by easing the control on the gene-network, allowing gene expression levels to fluctuate between cells, eventually leading to the emergence of a phenotype better adapted to the unforeseen stress [91, 92]. In contrast, in the second framework, dysregulation results from the failure of the evolved stress response. Similarly to “throwing a wrench into the gears of the regulatory network”, dysregulation is the consequence of strong perturbations to the large network of gene-gene interactions and the subsequent chaotic state [83], as our recent model of disruption suggests [1]. Which of these two scenarios underlies the disrupted growth-arrest remains an open question. In both cases, the GMP-Cor metric should provide an invaluable tool to quantify its dysregulation.

The intricate relation between antibiotic persistence and growth-arrest suggests that our findings should advance the general understanding of antibiotic persistence. While the search for persistence-enabling genetic pathways may be relevant in specific cases, our results suggest that tolerance and persistence can result from dysregulation under acute stress leading to Dis-Arrest, prior to a lethal antibiotic treatment such as ampicillin. While Dis-Arrest enhances survival under a growth-targeting antibiotic, it is not a straightforwardly advantageous state. In the same cells we measure both enhanced survival to ampicillin (Fig. 5B) and markedly increased sensitivity to 1% SDS (Fig. 5H). These two properties are not correlated, and there is no reason they should be. Tolerance to ampicillin arises from the absence of growth, whereas SDS acts on the envelope irrespective of growth state and reports envelope integrity. A cell can therefore be simultaneously tolerant to a growth-targeting antibiotic and have a compromised membrane. Persisters do not need to excel at both, and the expectation that they should stems from studies in which tolerance or persistence was attained from a mechanism other than dysregulation[93, 94].

This understanding implies that screens designed to pinpoint “persistence genes” may yield irreproducible hits whenever persistence arises from broad dysregulation. This is because the physiological behavior stems from a global property of the gene network, rather than of specific nodes. In these conditions, many gene deletions may enhance antibiotic persistence by increasing dysregulation, while deletions that restore regulatory balance are far less likely. Consequently, CRISPR or Tn-seq approaches seeking knockouts that decrease persistence will rarely identify universal targets, as suppression of persistence may strongly depend on how the underlying dysregulation was attained, rather than on a true universal “persistence gene”. We do not claim that persistence always arises from global dysregulation. Two accepted hypotheses remain, induction of defined pathways[33] versus a global property of the gene network[95], with evidence supporting both sides. Our results indicate that in the Dis-Arrest conditions studied here the evidence supports the latter, and identifies dysregulation as the specific form of broad response. In contrast, Reg-Arrest cells do activate the general starvation response known to result in broad tolerance to various stresses, as long as they remain in starvation conditions (Fig. 1A).

The results of this study suggest that the resilience of persistent bacteria to certain antibiotic treatments may be accompanied by key vulnerabilities, due to the general loss of regulation. Our initial findings indicated that increased membrane permeability may represent one such vulnerability to be further exploited, for example by searching for drug combinations against persistence [96]. Practically, we believe our results can be used as a disrupted state biomarker. Because dysregulated populations are enriched for long-lag, tolerant cells, a low GMP-Cor flags treatment regimens or strain backgrounds likely to generate persisters, and may identify conditions, including in the host, in which the sensitivity of disrupted cells to membrane-targeting agents could be exploited[2]. Moreover, we believe that this framework extends beyond bacterial antibiotic persistence. The focus on global dysregulation, and the method that we proposed to measure it, should be relevant also to other biological systems under acute stress, such as persister cancer cells under chemotherapy [97, 98, 99, 100]. We tested the validity of GMP-Cor on cancer single-cell data[3], and showed that it captures the difference in correlation levels between proliferating and arrested cells under anti-cancer treatment (supp. note 6, Extended Data Fig. S15). Further extending this analysis with physiological measurements, may reveal new key vulnerabilities of disrupted cancer cells.

## 4 Methods

### Bacterial strains and growth conditions

The SHX induced growth-arrest and natural starvation experiments were performed on *E. coli* KLYR [1]. This strain is characterized by a relaxed stringent response due to mutations in the *relA1* [101] and *spoT1* [102] genes. This relaxed stringent response is what makes the arrest induced by SHX disruptive. Importantly, the genetic background is not the source of the phenotype - the same strain administered SHX gradually, or allowed to enter stationary phase naturally, produces a regulated arrest with narrow lag-time distributions[1] (Extended Data Fig. S16). Furthermore, Dis-Arrest can equally be produced by stresses unrelated to the stringent response, such as chloramphenicol or sodium azide [1]. Bacteria were diluted 1:10^6^ from a frozen aliquot into fresh medium (M9, Sigma Aldrich) supplemented with 0.1% Casamino acids, MgSO_4_ (120µg/ml), vitamin B1 (0.5µg/ml) and kanamycin (30 µg/ml, Sigma). Bacteria were grown at 32° C with shaking. To achieve Reg-Arrest, cultures were allowed to grow in the same conditions followed by a natural depletion of nutrients. Reg-Arrest samples were collected 24 h after entry into stationary phase, defined at OD_600_ ≈0.65, ∼24 h after inoculation. To induce Dis-Arrest, cells were grown to exponential phase (∼13 generations from the initial inoculation, when cells reach ∼ 10^7^cells/ml, OD_600_ ≈0.1), following exposure to SHX at a concentration of 0.5mg/ml. Experiments involving SHX-induced Dis-Arrest were performed following a 24 hour exposure to SHX, unless stated otherwise.

VapC induced growth-arrest experiments were performed on *E. coli* MG1655 intRCmCH strain with a chromosomal insertion of the *vapC* and the mCherry genes after the Tet promoter, in proximity to constitutive expression of the tet repressor [73]. This allows induction of the VapC protein with anhydrotetracycline (aTc). For the Reg-Arrest control in this experiment, we used the MG1655 intRmCH strain, without the *vapC* gene. We verified that the induction of VapC does not introduce cell-to-cell variability by observing similar coefficient of variations (CV) for both strains (Extended Data Fig. S13B). In this experiment, bacteria were grown at 37*^◦^*C in M9 medium (Sigma Aldrich), supplemented with 0.1% casamino acids, MgSO_4_ (120µg/ml), vitamin B1 (0.5µg/ml) and 7.5µg/ml kanamycin. Cells were either diluted 1:10^8^ from a frozen aliquot into fresh medium or grown overnight in LB medium, washed twice (3 min, 5000g), and resuspended and diluted 1:100 in M9 medium, depending on the experiment. When the bacteria reached OD_600_ ≈ 0.2, aTc (100µg/ml stock in double-distilled water (DDW)) was added to the medium to a final concentration of 25ng/ml, inducing growth-arrest on the intRCmCH strain but allowing uninterrupted growth of the control strain, which eventually reaches growth-arrest due to a natural depletion of nutrients. In the single cell RNA-seq experiment, MG1655 intRCmCH strain includes an additional plasmid with a GFP reporter (S*GFP) under P_lac_ promoter. This strain was grown as explained above, but with 7.5µg/ml kanamycin and 100 µg/ml ampicillin. GFP was induced with 25µg/ml IPTG 1 hour after induction of *vapC* with aTc. For RNA-seq experiments, bacteria were grown in a liquid volume of 60 ml contained in a 250 ml Erlenmeyer flask. For all other experiments, bacteria were grown in 96-well plates with 180-200µl total volume in each well. Plates were incubated in a plate reader (Tecan Infinite M200 Pro) with orbital shaking at 270 rpm (2.5mm amplitude). We note that oxygenation influences outcomes of prolonged growth-arrest so culture volumes and shaking are important factors.

### Lag time measurements using ScanLag

A sample of the culture was diluted and plated on M9 agar plates supplemented with 1% casamino acids, MgSO_4_ (120µg/ml), vitamin B1 (0.5µg/ml) and kanamycin (15 µg/ml, Sigma) (for the S*GFP plasmid strain 100µg/ml ampicillin was added). Plates were placed on an automated scanner setup (ScanLag [71]) in an incubator at 37*^◦^* C and imaged every 15 minutes. The images were analyzed using ScanLag software, and the appearance time of each colony was measured (code is available at https://github.com/oferfrid/NQBMatlab/tree/V16). The median appearance time of the exponential control from each experiment is subtracted from raw colony appearance times to result in net lag times, as reported previously and validated by microscopy at the single cell level both on solid and liquid media[1]. We note that treatments are applied in liquid culture whereas lag times are read out on solid medium. The comparison is internally controlled, every condition is treated in liquid and plated identically, so a systematic liquid-to-solid effect cannot generate the differences between conditions.

### Single cell RNA sequencing

Samples were grown as explained in *Bacterial strains and growth conditions*. We collected 1ml from each culture following fixation with 1% formaldehyde (67µl of 16% stock was added to the sample). Samples were then incubated at room temperature for 30 minutes and spun down (5 minutes, 6000 rcf at room temperature). The supernatant was removed, and the pellet was resuspended in 1ml of 0.02X SSC (Invitrogen™). Samples were then spun down again (3 minutes, 6000 rcf at room temperature), the supernatant was removed, and the pellet resuspended in 350µl MAAM (4 parts methanol to 1 part acetic acid). Samples were kept in −20*^◦^*C until processed for scRNAseq. The single cell experiments were performed using ProBac-seq [55], a droplet based technique that was designed specifically for prokaryotic cells (see illustration in Fig. 1F). Probac-seq utilizes libraries of pre-synthesised DNA probes, thus removing the rRNA depletion step necessary in most bacterial single cell techniques available today. This is an advantage when studying starved bacteria, which suffer from even lower RNA counts than growing bacteria, where the signal could potentially be lost to residual rRNA sequencing. Samples in this study were processed using a protocol described elsewhere [55, 103], with minor additions: one probe against newly synthesized (unprocessed) rRNA targeting the 5’ premature 16S region, one probe for mature rRNA targeting the 16S ribosome, as well as probes targeting the highly abundant ncRNA target tmRNA and four probes against *yfp*, *mCherry*, *vapC*, *tetR*. The probes contain unique molecular identifiers (UMI) to enable PCR duplication error correction. After mapping the raw reads to the *E. coli* genome, we construct a cell-gene matrix that contains the amount of reads that were mapped to a specific gene for every unique cell barcode.

### Mapping of the single cell RNA-seq reads

Raw FASTQ files were processed with Cell Ranger 7.0.0 (10x Genomics) [104] using default parameters against a custom *E. coli* reference built with cellranger mkref (custom FASTA files are provided) [105]. Slight modifications to the original pipeline were made to adapt to the ProBac-seq [55] data. First, the 16nt 10x UMI from R1 and the 12nt probe UMI from R2 were stitched together using cutadapt [106]. Sequential adapters and unwanted sequences were trimmed, truncating R2 at 54nt and R1 at 28nt. Finally, count data was produced using cellranger count with –chemistry=SC3Pv3 (10x chemistry) and –transcriptome set to our custom genome.

### Initial processing of the single cell data

To analyze the count matrix, we must filter the barcodes that contain real cells from the barcodes that represent empty droplets with ambient RNAs. This is done by analyzing the Unique-Molecular-Identifier (UMI) count rank plot which depicts the total amount of reads for each cell barcode. We take a lower threshold of UMI counts that results in ∼ 1000 cells in the lower coverage dataset for each condition and apply the same threshold to each dataset, randomly selecting 1000 barcodes above this threshold. This way we mitigate the effect of different sample depth within the same condition and fix the matrix dimensions. For the correlation analysis, we take the top 2000 genes ranked by Fano factor over cells. GMP-Cor is an extensive quantity: it is a sum over the eigenvalues of a spectrum whose total mass equals the number of genes, so its magnitude scales linearly with the number of genes analyzed, and is also influenced by the dimensionality to sample size ratio 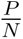. We show how GMP-Cor is influenced by sub-sampling cells and how it scales with gene number in Extended Data Fig. S6C-D. All comparisons in this study are therefore made between datasets of matched gene dimension, as fixed by this filtering step. For the analysis we omit the 16S rRNA probe that was introduced for cell calling as well as plasmid integrated genes to avoid the noise related to copy number. For calculating the distribution of eigenvalues of the correlation matrix we also normalize the total reads per cell, 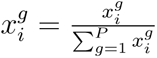, where i and g denote the cell and gene indices respectively. Further, we calculate the z-score of each gene 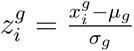. For this normalized matrix we calculate the gene correlation matrix 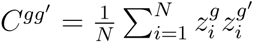 and its eigenvalues - the principal components. For UMAP, clustering and marker gene analysis, we use the scanpy python library (version 1.9.8) for scRNA-seq analysis [107]. We filter the cell barcodes as explained previously, normalize using scanpy’s normalize total and log transform the data. We perform PCA analysis on the concatenated dataset and then run the nearest neighbor algorithm followed by UMAP embedding. We run nearest neighbors with n neighbors=40 and n pcs=40 and UMAP with min distance=0.3. Clustering is performed on the gene expression data using the Leiden algorithm [108] (resolution=0.4/0.3 in Fig. 1 and Fig. 5 respectively) and marker genes are identified by performing the Wilcoxon rank test.

### Applying tools from Random Matrix Theory to measure correlations in high-dimensional data

To illustrate the ways spurious correlations may impede the measurement of correlations in high-dimensional data, we draw an example from the Wishart random matrix ensemble [109]. In this ensemble, the correlation matrix is constructed as the average of *N* contributions:

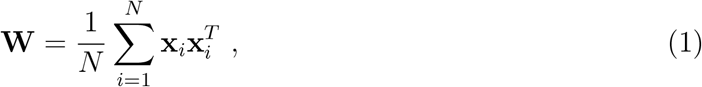

where 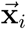 is a *P*-dimensional normally distributed random vector with independent components, satisfying 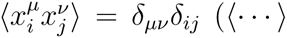 denotes averaging over an infinite ensemble). To associate this framework with the experimental data, *N* should be interpreted as the number of cells, *P* is the number of genes, and 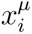 represents the expression level of gene *µ* in the i-th cell. At first glance, this construction should result in an absence of correlations between different genes. However, the exact eigenvalue distribution of this matrix follows the celebrated Marchenko-Pastur (MP) distribution [70]:

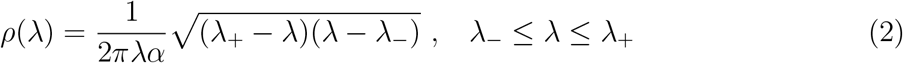

with 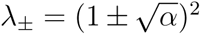 and *α* = *P*/*N*. The finite width of this distribution, 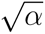, for any finite size ensemble (*N* < ∞) results in the emergence of finite correlations, even though all vector entries are statistically independent by construction. Furthermore, in typical experimental settings where the dimensionality-to-sample size ratio, *α*, is on the order of one, substantial spurious correlations are introduced. Note that when *N* ≥ *P*, the correlation matrix is full rank, and all its eigenvalues must be positive by definition. However, when *N* < *P* (i.e. *α* > 1), the correlation matrix has rank *N* resulting in *P* − *N* zero eigenvalues. These zero eigenvalues contribute a δ-function at the origin in *ρ*(λ). Since this δ-function does not provide meaningful information, it will be omitted from further analysis.

The example above demonstrates that uncovering genuine correlations in experimental data necessitates a method capable of effectively removing spurious correlations introduced by finite-size effects. To address this, our approach assumes that, as a first approximation, genes are correlated, but to an observer who does not know about the details of the gene-network - these correlations appear random. In this framework, the entries of the vectors 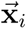 are sampled from a multivariate Gaussian ensemble with a covariance matrix defined as:

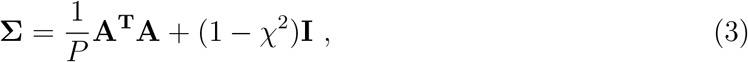

where *A* is a *P* ×*P* random matrix with independent entries, 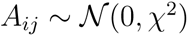. The parameter χ ∈ [0, 1) tunes the level of dependencies between random variables in the true covariance matrix, with χ = 0 recovering the standard Wishart ensemble. The eigenvalue distribution of the resulting ensemble of correlation matrices, referred to as the Generalized MP (GMP) distribution, can be calculated exactly (see Supp. note 1 for a detailed derivation). The GMP distribution gives us the added effect of genuine correlations in the data and can be used to separate this signal from the spurious correlations.

### Quantifying the global correlations in the data

To quantify the level of correlations in the data, we define GMP-Cor as the sum of eigenvalue differences from the maximal scrambled eigenvalue, counting positive values only:

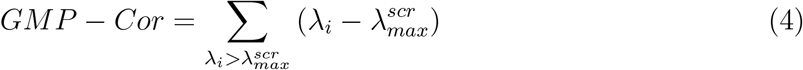

This value takes into account both the quantity and absolute value of the large eigenvalues, which we interpret as the correlation signal. Normalizing each eigenvalue by the maximum scrambled eigenvalue ensures that if an eigenvalue crossed this threshold just by an infinitesimal amount, possibly due to sampling noise, the value of GMP-Cor won’t jump discontinuously.

### GMP-Cor permutation test

To infer the statistical significance of GMP-Cor, we perform a permutation test on the maximal scrambled eigenvalue. To do this, we rerun the scrambling procedure 2000 times and record the value of 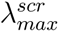 each time. We use this data for two purposes:

1. We calculate an empirical p-value for the significance of the difference between the original and scrambled eigenvalue spectra. This is defined as

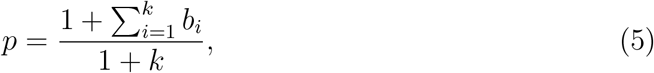 where 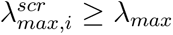 and *b_i_* = 0 otherwise. k is the number of permutations.
2. We calculate σ (standard deviation) of 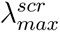 and then define the confidence interval of GMP-Cor as 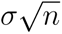, where n is the number of eigenvalues that cross the 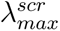 threshold and are included in the GMP-Cor summation.

### Bulk RNA sequencing

Bulk RNA-seq data comparing Dis-Arrest, Reg-Arrest and Exponential growth conditions was obtained from Rotem et al. [2]. Our analysis of this data is presented in Fig. 4A-B. The experiment investigating different time points in SHX treatment was performed as follows: Cells were grown to Dis-Arrest condition as described in *“Bacterial strains and growth conditions”*. Samples of 1.8ml were collected at the specified times after exposure to SHX. Each sample was spun down for 5 minutes at 5000g, 4*^◦^*C. After spin down, the excess fluid was discarded and the pellet resuspended in 50µl TE buffer (10mM Tris-HC1 pH 7.5, 1mM EDTA), inserted into a precooled Eppendorf with 5µl of 9mg/ml Lysozyme in RNase-free water (Sigma), briefly submerged in liquid nitrogen, and stored at −80*^◦^*C.

The frozen samples were subjected to two cycles of thawing at 37*^◦^*C and refreezing in liquid nitrogen. Next, the samples were resuspended thoroughly to homogenization with 1 ml TriReagent and incubated for 10 min at room temperature. Chloroform (200 µl) was added, and the tubes content was mixed by inversion for 15s. The samples were incubated for 10 min at room temperature, centrifuged (14,000g, 10 min at 4*^◦^*C) and the upper phase was collected and transferred into new Eppendorf tubes. For RNA precipitation, 500 µl isopropanol was added, the tube contents were mixed thoroughly by inversion and incubated for 10 min at room temperature. The tubes were centrifuged (14,000g, 15 min at 4*^◦^*C) and the supernatant was discarded. The pellets were washed twice by addition of 1 mL of freshly made 75% ethanol, followed by centrifugation (14,000g for 5 min at 4*^◦^*C) and removal of the supernatant. Next, samples were spun down for 3 min at room temperature, and the excess fluid was discarded. Pellets were dried by leaving the tubes open for 10-15 min at room temperature, and then re-suspended in 50µl RNase-free water and stored at −80*^◦^*C. RNA quality and concentration was assessed by Nanodrop (ThermoFisher Scientific) and by Bioanalyzer (Agilent).

rRNA depletion, library preparation and RNA-seq was done by the center for genomic technologies, the Institute of Life Sciences, the Hebrew university of Jerusalem. rRNA depletion was done using NEBNext rRNA depletion kit for bacteria. Library preparation was done using NEBNext ultra II directional RNA library prep kit. RNA-seq was done using Illumina NextSeq 500.

### Bulk RNA-seq data analysis

Read trimming was done using Cutadapt [106] (With parameters: -e 0.2 -m 15 -O 1 -q 20). Mapping was done using bowtie2 [110] (With parameters: –local –very-sensitive-local). The raw reads were mapped to the genome available at CP008801.1. Expression counting was done using htseq-count [111] (with parameters: stranded option, –nonunique none). For the dynamics under SHX, the bulk transcriptomics raw count data was normalized using ERCC spike-ins [112] by calculating the median level of ERCC spike-in levels relative to their known concentration (mix 1, concentration datasheet available from ThermoFisher), and using this value as a scale factor between samples. This technique is used to reduce the high technical variability of total counts between samples that often occurs in RNA-sequencing experiments.

### GO enrichment analysis of the bulk RNA-seq data

The gene ontology (GO) enrichment analysis was performed on the set of differentially expressed genes (DEGs) obtained from the DESeq2 algorithm [113], and implemented using the Python package PyDESeq2 (version 0.4.4) [114]. The log_2_ fold change (LFC) results for both the Dis-Arrest and Reg-Arrest conditions are compared to the exponential condition as the control group. Statistical significance of DEGs is assessed by DESeq2, based on 5 samples for each condition. To test if DESeq2 normalization creates a bias for conditions with a different amount of total transcripts, we also performed ERCC spike-in normalization (see above) and show that the results are qualitatively the same for both normalization schemes (Supp. Note 5, Extended Data Fig. S12). While both conditions reduce most transcripts compared to exponential growth, the study set of genes for the GO enrichment test was defined as the 1000 most prominently reduced genes (according to the LFC) that also passed the threshold of p*_adj_* < 0.01. This ensures that we compare GO enrichment results correctly from study sets of the same size. Enrichment analysis was performed using the Python package GOATOOLS (version 1.6.4) [115]. This algorithm implements *Fisher’s exact test* [116] on the set of DEG’s compared to the background set, and then performs the *Benjamini-Hochberg method* [117] to adjust the resulting p-values for multiple tests. A threshold for selecting significant GO terms was set at p*_adj_* < 0.05. For the time dependent experiment (Fig. 4C), one sample was collected for each of the 8 time points in SHX stress, in addition to an initial time t_0_. The bulk data was normalized by ERCC spike ins, and the LFC was calculated in comparison to the initial time point (t_0_). The 400 most downregulated DEG’s (according to the LFC) were considered as the study set, and enriched GO terms were calculated based on this set. To mitigate the variability caused by a lack of repetitions at each time-point, marginally enriched terms were omitted from this analysis, and only GO terms that were significantly enriched in all of the time-points were considered.

### Antibiotic killing assay

VapC induced growth arrest, as well as Reg-Arrest control cells were grown as described in *Bacterial strains and growth conditions*. At 24h after aTc addition (initiation of VapC growth-arrest), cultures were washed, diluted 1:50 into fresh growth medium and incubated for 2h. Ampicillin was then added to a final concentration of 100µg/ml (from a 100mg/ml stock in DDW). At the indicated time-points, viable cell counts were determined using the Most Probable Number (MPN) method. For each culture and time-point, 96-well plates were prepared with LB medium and 12.5µg/ml kanamycin. Cells were distributed in each plate with an initial dilution of 1:50 in the first column, followed by serial dilutions of 1:6 for each subsequent column, with the 8 rows serving as technical replicates. Plates were then incubated at 37*^◦^*C and wells were scored for growth.

### SDS assay

VapC induced growth arrest, as well as Reg-Arrest control cells were grown as specified in *Bacterial strains and growth conditions*. 10% SDS stock solution was prepared by adding SDS powder to DDW at a mass ratio of 1:10. The solution was further filtered for sterilization. SDS stock solution was diluted 1:10 in 200µl wells to achieve a final concentration of 1%. Cells were then incubated with the SDS solution and OD_600_ measurements were performed using a Tecan™Infinite M200 pro plate reader.

### Microscopy imaging

Microscopy was done using a Nikon Ti2E inverted fluorescence microscope system, ×100 NA (numerical aperture) 1.45 oil objective with phase-contrast, automated stage and shutters, and automated focus hardware (Perfect Focus System - PFS). Temperature was controlled using a cage incubator (Okolab). The control of the microscope, stage, shutters, camera, PFS, and image acquisition were done using Micro-Manager open-source microscopy software. Images were acquired with a sCMOS digital camera (Orca Flash 4.0v3, Hamamatsu). YFP imaging: Spectra X (Lumencor), EX filter 500/24 EM 542/27 (Semrock). Red fluorescence imaging (used for mCherry measurements): Spectra X (Lumencor), EX filter 562/40 EM 640/75 (Semrock).

### Microscopy image analysis

Phase-contrast frames were segmented with a pretrained Cellpose-SAM model running on GPU. Images were normalized with inversion before evaluation, one binary mask layer was written per frame. Fluorescence background was modeled per image as a tilted plane fit to pixels outside the cell mask. The image was detrended by subtracting the plane. An object was marked as a cell only if (i) its median fluorescence exceeded the background median by ≥ 2σ, and (ii) its median phase intensity was below the global phase background. For the accepted cells, area was measured as the mask pixel count, length was measured as the major axis of a fitted ellipse, and mCherry intensity as the per-cell median of the background-subtracted mCherry image. Pixel counts were converted to units of length using a calibrated measurement of 15.15px to 1µm.

### Microfluidics time-lapse analysis

Observations of bacteria in microfluidic mother-machine devices were made as previously described [1]. Movies were acquired at 10 min/frame with the drug added at frame 18; 50 frames were analyzed. Phase-contrast images were processed with DeLTA’s pretrained mother-machine U-Nets for trench detection, cell segmentation and lineage tracking. Tracks were filtered in five stages. (1) YFP test: median background subtracted YFP > 2σ per frame, with a track retained if ≥ 50% of its frames were YFP-positive. (2) A frame was excluded if the total mask cell area jumped discontinuously from the previous frame. (3) Spurious division threading: a mother and a single daughter appearing at the mother’s terminal frame were merged into one track. (4) Post-drug births were retained only if the mother showed elongation before the event and the birth length was < 0.65× the mother’s pre-split length. (5) Tracks were required to span ≥ 5 frames. Divisions were validated by five tests: each daughter’s birth length < 0.65× the mother’s pre-split length; the products’ summed length within ±20% of the mother’s pre-split length; every product observed ≥ 3 frames; the mother elongated ≥ 1.4x since its birth or previous validated division; and, a measurable increase in cell boundary length calculated on the YFP intensity at the mask boundary. Of 405 putative divisions, 62 passed all tests. To assess cell fate, integrated growth G was computed on a 3-frame centered rolling median of log elongation. With D the number of validated divisions in the window, cells were classified as halted (G < g*_l_*and D = 0), growing (G ≥ g*_h_* or D≥1) or ambiguous. Thresholds were calibrated to fit known growing or halted cells. Halt time was determined as the first frame at or after drug addition where the elongation rate per frame µ = Δ ln L remained below 0.03 for 5 consecutive observed frames, and the cell had grown earlier in the movie. Halt times are reported for cells classified as halted. **Software**: Cellpose 4.2.1 (Cellpose-SAM), PyTorch 2.11.0 (CUDA 12.8), NumPy 2.4.6, SciPy 1.18.0, scikit-image 0.26.0, tifffile 2026.6.1, OpenCV 4.13.0, DeLTA 2.0.7, TensorFlow 2.10.1

### Statistical analysis and tests

All statistical tests were performed using the python library scipy.stats. For evaluating significance on the GMP-Cor metric and enrichment score of GO analysis we used the 2 sided Mann-Whitney U-test. To obtain confidence intervals for Scan-lag data (Fig. 1E) and enrichment score over time (Fig. 4C), we calculated the t-score that corresponds with a 68% confidence interval from a t-distribution with *n_samples_* − 1 degrees of freedom (in python: scipy.stats.t.ppf(0.84, df=n-1)). Then the error is given by 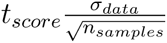.

## Acknowledgements

We thank A. Erez and R. Faigenbaum-Romm for fruitful discussions and constructive comments on the manuscript, Tomer Naor for assistance with preprocessing the scRNA-seq data, and an anonymous reviewer for insightful comments. This work was supported by the European Research Council (advanced grant no. 101054653), The Israel Science Foundation (grant no. 597/20) and the Minerva Center for Stochastic Decision making in Microorgan-isms. A.G. acknowledges the support of the ADAMS fellowships program and the Milner foundation.

## Author contributions

A.G, O.A and N.Q.B conceived the project. A.R, I.R, A.Z.R and A.G performed experiments. A.G performed data analysis and developed software. A.G, O.A and N.Q.B wrote the manuscript with input from all co-authors.

## Data availability

All raw sequencing data, including mapped reads were submitted to ArrayExpress under accession number E-MTAB-15651/ E-MTAB-15652. Additional data, including Scanlag and microplate reader experiments are available in the github repository: https://github.com/alongutf/dysregulated-persistence

## Code availability

All code developed for data analysis and generation of figures is available in the github repository https://github.com/alongutf/dysregulated-persistence.

