## Supplementary Notes for "GMP-Cor: Quantifying Transcriptional Dysregulation in Antibiotic Tolerance and Persistence"

#### 1 Analytical calculation of the generalized Marchenko-Pastur distribution

In this note, we provide the detailed calculation of the eigenvalue distribution for random correlation matrices (Wishart matrices) with a non-identity true covariance. The calculation roughly follows the known replica symmetric solution of the original Marchenko-Pastur (MP) distribution, but does not assume independent and identically distributed (iid) variables in the construction of the Wishart matrix.

We first define the Wishart matrix,

$$\mathbf{W} = \frac{1}{N}\sum_{i=1}^{N} \mathbf{x}_i \mathbf{x}_i^T, \tag{1}$$

where $\mathbf{x}_i$ is a $P$-dimensional random vector drawn from a multivariate Gaussian distribution $\mathbf{x} \sim \mathcal{N}(0, \Sigma)$. The probability distribution function (pdf) is given by:

$$p(\mathbf{x}) = \frac{1}{\sqrt{(2\pi)^P |\det \mathbf{\Sigma}|}} \exp\left(-\frac{1}{2}\mathbf{x^T \Sigma^{-1} x}\right). \tag{2}$$

Next, we write down the expression for the density of eigenvalues:

$$\rho(\lambda) = -\lim_{N\to\infty}\lim_{\epsilon\to 0^+} \Im \frac{2}{\pi P}\frac{\partial}{\partial\lambda}\left[\log\left(\det\left[(\lambda - i\epsilon)\,\mathbf{I} - \mathbf{W}\right]\right)^{-\frac{1}{2}}\right], \tag{3}$$

where $\Im$ is the imaginary part.

Defining $\mathbf{M} = (\lambda - i\epsilon)\,\mathbf{I} - \mathbf{W}$ we rewrite: $\log\det\mathbf{M}^{-\frac{1}{2}} = \lim_{n\to 0}\frac{\left(\det\mathbf{M}^{-\frac{1}{2}}\right)^n - 1}{n}$.

This limit can also be written as $\lim_{n\to 0}\frac{\partial}{\partial n}Z^n$, where $Z = \det\mathbf{M}^{-\frac{1}{2}}$.

Now, we write the determinant as a product of Gaussian integrals:

$$Z^n = \int_{-\infty}^{\infty}\prod_{a=1}^{n} d\mathbf{u}^a \exp\left(-\frac{\lambda - i\epsilon}{2}\sum_a \mathbf{u}^{aT}\mathbf{u}^a + \frac{1}{2}\sum_a \mathbf{u}^{aT}\mathbf{W}\mathbf{u}^a\right) \tag{4}$$

The second term in the exponent can be rewritten as: $\frac{1}{2}\sum_a \mathbf{u}^{aT}\mathbf{W}\mathbf{u}^a = \frac{1}{2}\sum_a\sum_i \left(v_i^a\right)^2$, where $v_i^a = \frac{1}{\sqrt{N}}\mathbf{x}_i^T \cdot \mathbf{u}^a$.

The statistics of $v$ are given by $\langle v_i^a v_j^b \rangle = \delta_{ij} Q^{ab}$, where $\langle \cdot \rangle$ denotes averaging over the disorder and $Q^{ab} = \frac{1}{N}\sum_{\mu,\nu} u_\mu^a u_\nu^b \langle x_\mu x_\nu \rangle = \frac{1}{N}\sum_{\mu,\nu} u_\mu^a u_\nu^b \Sigma_{\mu\nu}$.
We can work in the eigenbasis of $\Sigma$ without affecting the results.

$$Q^{ab} = \frac{1}{N}\sum_\mu u_\mu^a u_\mu^b \sigma_\mu^2 \tag{5}$$

Returning to the second term in the exponent of eq. 4, we have:

$$\begin{aligned} \exp \frac{N}{2}\sum_a (v_i^a)^2 &= \left( \int \frac{dv^a}{(2\pi)^{\frac{n}{2}}\sqrt{\det \mathbf{Q}}} \exp\left( -\frac{1}{2}\sum_{ab} v^a \left[ Q^{ab^{-1}} - I^{ab} \right] v^b \right) \right)^N \\ &= \exp -\frac{N}{2}\left( \log\det\left[ \mathbf{Q}^{-1} - \mathbf{I} \right] + \log\det \mathbf{Q} \right) = \exp\left( -\frac{N}{2}\log\det\left( \mathbf{I} - \mathbf{Q} \right) \right) \end{aligned}$$

Next, integrating over $u^a$ and imposing the constraint defined in eq. 5 using an integration variable $\hat{Q}$ yields,

$$\langle Z_n \rangle = \int d\hat{Q}^{ab} \int dQ^{ab} \exp\left( -i\frac{N}{2}\sum_{ab} \hat{Q}^{ab} Q^{ab} - \frac{P}{2}\log\det\left[ (\lambda - i\epsilon)\mathbf{I} - i\hat{\mathbf{Q}}\Sigma \right] - \frac{N}{2}\log\det\left[ \mathbf{I} - \mathbf{Q} \right] \right) \tag{6}$$

The replica symmetric ansatz implies that there is no overlap between replicas, therefore we have:

$$\begin{aligned} Q^{ab} &= q\delta^{ab} \\ \hat{Q}^{ab} &= -i\hat{q}\delta^{ab} \end{aligned}$$

$$\langle Z_n \rangle = \int d\hat{q} \int dq \exp\left( -\frac{Nn}{2} f(q, \hat{q}) \right) \tag{7}$$

$$f(q, \hat{q}) = q\hat{q} + \frac{1}{N}\sum_\mu \log\left( \lambda - i\epsilon - \hat{q}\sigma_\mu^2 \right) + \log(1 - q) \tag{8}$$

The saddle-point equations for $q$ and $\hat{q}$ yield,

$$\hat{q} = \frac{1}{1 - q} \tag{9}$$

$$q = \frac{1}{N}\sum_\mu \frac{\sigma_\mu^2}{\lambda - i\epsilon - (1 - q)^{-1}\sigma_\mu^2} \tag{10}$$

From now on we denote the eigenvalues of the true covariance matrix $\sigma_\mu^2$ as $x$. We can solve the second equation if we have a closed form for $\rho(x)$. Given our definition of $\boldsymbol{\Sigma} = \frac{1}{P}\mathbf{A}^\mathbf{T}\mathbf{A} + (1 - \chi^2)\mathbf{I}$, we know that the first term gives us the original MP distribution with $\frac{P}{N} := \alpha = 1$ (defined in Methods), and the second term is proportional to the identity and therefore gives a simple shift of the eigenvalues $x \to x + 1 - \chi^2$.

$$\rho(x) = \frac{1}{2\pi\chi^2(x - 1 + \chi^2)}\sqrt{(5\chi^2 + 1 - x)(x - 1 + \chi^2)} \tag{11}$$

Converting the sum in eq. 10 to an integral over the eigenvalues we replace, $\sum_\mu \to P \int dx \rho(x)$. Calculating the integral yields,

$$q = \alpha(q-1)\left(1 - \frac{\lambda(q-1)\left(\sqrt{\frac{\lambda(q-1)+3\chi^2+1}{\lambda(q-1)-\chi^2+1}}-1\right)}{2\chi^2}\right) \quad (12)$$

This equation reduces to a fourth degree polynomial for $q$ and its closed form solution is too long to be listed here, but can be obtained with Wolfram Mathematica (see Mathematica notebook in the github repository). There are 2 complex conjugated pairs of solutions, though one of these pairs is un-physical. We denote the physical solution with the positive imaginary part as $q(\alpha, \chi, \lambda)$. Substituting the results for $q$ and $\hat{q}$ in eq. 7 we get:

$$\langle Z^n \rangle = \exp\left(\frac{-Nn}{2}\left[\frac{q}{1-q} + \alpha \int dx \rho(x) \log\left(\lambda - i\epsilon - (1-q)^{-1}x\right) + \log(1-q)\right]\right), \quad (13)$$

Using eq. 3, we get a closed form of the eigenvalue distribution,

$$\rho(\lambda) = -\lim_{N\to\infty} \lim_{\epsilon\to 0^+} \lim_{n\to 0} \Im \frac{2}{\pi P} \frac{\partial}{\partial \lambda} \frac{\partial}{\partial n} \langle Z^n \rangle \quad (14)$$

$$= \Im \frac{1}{\pi} \int dx \frac{\rho(x)}{\lambda - (1 - q(\alpha, \chi, \lambda))^{-1} x}. \quad (15)$$

Finally, we obtain,

$$\rho(\lambda) = \Im \frac{(q(\lambda, \alpha, \chi) - 1)}{2\pi\chi^2}\left(\sqrt{\frac{\lambda(q(\lambda, \alpha, \chi) - 1) + 3\chi^2 + 1}{\lambda(q(\lambda, \alpha, \chi) - 1) - \chi^2 + 1}} - 1\right), \quad (16)$$

the results of which are plotted in Extended Data Fig. S1B. To build intuition for the results, consider a scatterplot of three-dimensional random Gaussian data. By adjusting the parameter $\chi$, we can compare cases of identically and independently distributed (iid) variables with those of non-iid variables. In the iid case, the eigenvectors of the correlation matrix—known as the Principal Components (PCs)—form a unit sphere (Extended Data Fig. S1**A**). In contrast, for non-iid variables, the PCs are stretched and contracted, spanning an ellipsoid (Extended Data Fig. S1**B**). While direct visualization of the data is impossible in high dimensions, the same pattern of stretching and contraction is reflected in the eigenvalue distributions predicted by the generalized MP distribution.

We next assess the validity of the analytical results. Although we do not provide a rigorous proof that the replica symmetry assumption holds or that the analytical continuation of $n \to 0$ is formally justified, our findings are supported by extensive testing with computer-generated data. These tests demonstrate excellent agreement with analytical predictions across a wide range of parameters. An example, using $\alpha = 0.5$ and $\chi = 0.7$, is shown in Extended Data Fig. S1**C**.

### 2 Synthetic Single-Cell RNA-seq Data Generator

#### Overview

To validate that the GMP-Cor metric faithfully reports gene-gene correlation strength under realistic noise conditions, we constructed a synthetic scRNA-seq data generator. The generator reproduces three key features of real single-cell data: (i) a structured, non-random gene-gene correlation matrix. (ii) heavy-tailed, overdispersed counts consistent with a negative-binomial model. (iii) expression-dependent dropout sparsity. The two-stage design - first fixing the correlation structure, then sampling counts through a copula (linking) distribution decouples the ground-truth correlation parameter from count-level noise, enabling controlled calibration of GMP-Cor across a range of correlation strengths $\chi$.

#### Stage 1: Constructing the correlation matrix

**Factor-model architecture.** The ground-truth gene-gene correlation matrix $R \in \mathbb{R}^{P \times P}$ is built from a loading matrix $A$ that encodes two levels of shared variance: *cluster factors* group nearby genes into co-expressed modules, and a *global hub factor* links a random subset of cluster hubs to create long-range correlations between modules. This topology mimics transcriptional regulation at both the operon/regulon level (clusters) and the master-regulator level (global hub).

**Loading matrix construction.** Let $P$ be the number of genes and $\chi \in (0, 1)$ the shared-variance fraction. The loading matrix is initialized as

$$A = \sqrt{1-\chi}\, I_n, \tag{17}$$

so that each gene's variance equals $1-\chi$. Cluster factors are appended as follows. Genes are partitioned into contiguous clusters whose sizes $s_k$ are drawn independently from a Pareto distribution:

$$s_k \sim \text{Pareto}(\text{shape}) + 1, \qquad k = 1, 2, \ldots, K, \tag{18}$$

until all $P$ genes are assigned. For cluster $k$ the shared loading of its member genes is

$$a_{ik} = \sqrt{w_k}, \quad w_k \sim \text{Uniform}(0.1\chi,\, \chi), \quad i \in \mathcal{C}_k, \tag{19}$$

and $a_{ik} = 0$ otherwise. A single global-hub column is then added: for each cluster hub $h_k$ (one gene chosen uniformly from $\mathcal{C}_k$), the hub weight is

$$a_{h_k,\,\text{hub}} = \begin{cases} \text{Uniform}(-\chi,\, \chi) & \text{with probability } p_{\text{hub}}, \\ 0 & \text{otherwise.} \end{cases} \tag{20}$$

The resulting loading matrix $A \in \mathbb{R}^{P \times (P+K+1)}$ satisfies $\tilde{C} = AA^\top$ is positive definite by construction.

**Normalization to a correlation matrix.** The positive-definite matrix $\tilde{C}$ is normalized to a correlation matrix by

$$R_{ij} = \frac{\tilde{C}_{ij}}{\sqrt{\tilde{C}_{ii}\,\tilde{C}_{jj}}}, \tag{21}$$

which preserves the off-diagonal structure while setting all diagonal entries to unity. The parameter $\chi$ directly controls the mean magnitude of the off-diagonal entries: larger $\chi$ produces stronger, more widespread correlations across genes.

#### Stage 2: Generating count data from latent Gaussian variables

**Latent multivariate normal sampling.** Given the correlation matrix $R$, we draw $N$ latent vectors, where $N$ represents the number of cells:

$$\mathbf{z}_c \sim \mathcal{N}(\mathbf{0},\, R), \qquad c = 1, \ldots, N. \tag{22}$$

These latent values encode the desired gene-gene correlation structure but have Gaussian marginals.

**CDF transformation.** We map latent values to the unit interval by applying the standard normal CDF gene-wise:

$$u_{ci} = \Phi(z_{ci}) \in (0, 1), \tag{23}$$

preserving all rank correlations. Where $\Phi$ is the Gaussian CDF. Because $\Phi$ is a monotone transformation, the Spearman correlations among genes are identical in the latent and uniform layers.

**Negative-binomial count marginals.** Gene counts are obtained by applying the negative-binomial (NB) quantile function to each uniform variate. For gene $i$, the mean expression level $\mu_i$ is drawn from an inverse-Gamma distribution,

$$\mu_i \sim \text{Inv-Gamma}(\kappa,\, \theta), \tag{24}$$

where shape $\kappa$ and scale $\theta$ are user-specified parameters ($\kappa = 2$, $\theta = 0.1$ in the representative simulation). This heavy-tailed prior reproduces the wide dynamic range of gene expression observed in real scRNA-seq data. The NB dispersion parameter is fixed at $r = 0.5$, a value that fits well with the single-cell data. The count for cell $c$ and gene $i$ is then

$$x_{ci} = F^{-1}_{\text{NB}(r,\, r/(r+\mu_i))}(u_{ci}), \tag{25}$$

where $F^{-1}_{\text{NB}}$ denotes the NB quantile (percent-point) function.

**Cell-level library-size variability.** To model differences in sequencing depth and cell size, each cell $c$ is assigned a multiplicative amplitude

$$\ell_c \sim \mathrm{LogNormal}(0.5,\, 1), \tag{26}$$

and the true count matrix is rescaled to $\tilde{x}_{ci} = \mathrm{round}(\ell_c \cdot x_{ci})$.

**Expression-dependent dropout.** Consistent with the technical dropout model widely used for scRNA-seq [1], a zero-inflation step masks observed counts as

$$y_{ci} = \begin{cases} \tilde{x}_{ci} & \text{with probability } 1 - e^{-\lambda \tilde{x}_{ci}}, \\ 0 & \text{with probability } e^{-\lambda \tilde{x}_{ci}}, \end{cases} \tag{27}$$

where $\lambda$ is the dropout rate parameter (default $\lambda = 1$). This formulation ensures that lowly expressed genes are preferentially dropped, reproducing the characteristic expression-detection correlation seen in real data. The simulation finally returns the observed (post-dropout) matrix $Y$ (Extended Data Fig. S2).

#### GMP-Cor calibration

The synthetic data generator is used to create a theoretical bound for the GMP-Cor value and to map the observable measure to a ground-truth correlation strength. We can achieve this by running a calibration sweep. A single correlation matrix $R_\chi$ is generated per $\chi$ value and held fixed across $n_{\mathrm{repeats}}$, so that the only source of variation between replicates is count-sampling stochasticity. GMP-Cor is computed for each replicate, yielding a median and standard deviation at each $\chi$ level. This calibration curve (Extended Data Fig. S4A) confirms that GMP-Cor increases monotonically with $\chi$ and is robust to count noise. We further compare GMP-Cor of the experimental data to the calibration curve and infer that the median regulated dataset translates to a correlation strength $\chi \approx 0.85$, while the median Dis-Arrest dataset translates to a correlation strength $\chi \approx 0.6$. This shows that while some of the Dis-Arrest samples produce a similar GMP-Cor to a completely random matrix, the count noise and sparsity screen most of the true correlation signal and the actual correlation strength is moderate.

#### Parameter choices and reproducibility

The correlation matrix seed and count-sampling seeds are stored separately, making it straightforward to re-run only the stochastic count layer while keeping the ground-truth correlation structure fixed. Default parameters used for the representative simulations shown in Extended Data Fig. S2 are summarized in the following table:

**Table 1:** Default parameters for the representative simulations.

| Parameter | Symbol | Value |
|---|---|---|
| Number of genes / cells | $P$ | 400 |
| Shared-variance fraction | $\chi$ | 0.9 |
| Pareto shape (cluster sizes) | $s$ | 1.5 |
| Hub-connectivity probability | $p_{\text{hub}}$ | 0.2 |
| Inv-Gamma shape | $\kappa$ | 2 |
| Inv-Gamma scale | $\theta$ | 0.1 |
| NB dispersion | $r$ | 0.5 |
| Library-size log-normal mean | $\mu_\ell$ | 0.5 |
| Library-size log-normal SD | $\sigma_\ell$ | 1.0 |
| Dropout rate | $\lambda$ | 1 |

### 3 Eigenvector analysis

GMP-Cor is a scalar that summarizes the overall gene-gene correlation strength of a sample from the eigenvalue density of its correlation matrix. A natural follow-up question is whether the eigenvectors associated with the largest eigenvalues can be used to recover which genes drive the coordinated structure, and how many genes participate in each coordinated mode. Here we analyze the leading eigenvectors of every scRNA-seq sample and show that the leading modes carry real and interpretable biology. However, the level of regulation is expressed in the magnitude of the eigenvalues, and not in the eigenvector loadings, which is why a single scalar that integrates most of the transcriptome (GMP-Cor), is the appropriate measure in our study.

#### Eigenvector loadings and gene coordination score

For each sample we diagonalized the gene-gene correlation matrix and retained not only the eigenvalues $\lambda_k$ but also the eigenvectors $\mathbf{v}^{(k)}$. The loading of gene $i$ in mode $k$ is the absolute amplitude $\left|v_i^{(k)}\right|$. We asked if we can recover what genes or programs contribute to the regulation or dysregulation we observe with GMP-Cor.

##### Results

1. We performed GO enrichment analysis for the top 100 genes in the 5 top modes for each sample (separating positive and negative loadings) and found significant enrichment of translation related GO terms in 12 of the 15 samples analyzed, spread across all conditions. This is a coherent, well-defined program, and it shows that the leading modes carry a biological signal. Other coherent programs we recovered from this analysis are chemotaxis and motility related terms (appearing only in exponential and early VapC samples), nucleotide biosynthesis (exponential cells), lipopolysaccharide biosynthesis

and stress response (appearing in both Reg-Arrest and Dis-Arrest samples). Two samples did not show any enrichment of GO terms: one Dis-Arrest sample and the 24h VapC sample.

2. In a few samples, we observe separate programs in the positive and negative loadings within the same mode. Genes with opposite signs in the same mode are anti-correlated and therefore represent an axis that separates cells between different expression patterns. In exponentially growing cells, we observe an axis that separates motility and nucleotide biosynthesis, and another axis that separates motility and translation. In one Dis-Arrest sample we observe a contrast between translation and lipopolysaccharide biosynthesis. Furthermore, we recognize a few RpoS-dependent general stress genes (although not enough to result in a significantly enriched GO term) that oppose the translation genes in the exponential cells, describing an axis running from high ribosome content to high stress-response. This is a real, interpretable biological signal obtained directly from analyzing the highest eigenvectors.

3. Genes transcribed from the same promoter appear together in the same mode, with the same sign. Our VapC strains carry a chromosomal integration of VapC and mCherry after the tet promoter. In the leading modes of these samples all these genes appear among the top-loading genes and carry the same sign. Because these genes are expected to be transcribed together, this acts as a positive control: the analysis groups co-expressed genes into the same mode.

#### Limitations

1. The eigenvectors do not distinguish Reg-Arrest from Dis-Arrest. The translation program appears in the leading mode of Dis-Arrest samples just as it does in Reg-Arrest samples and in exponentially growing cells. No other program appears consistently in one condition and not the other.

2. For nearly degenerate eigenvalues (which is the case of many of the top modes in Dis-Arrest), the corresponding eigenvector is not stable and can vary widely, by small changes in cell calling for example. This means that if 2 modes have a similar eigenvalue, it is impossible to assign unique genes to each of them and therefore it can be very difficult to interpret their biological meaning.

3. Importantly, bacterial scRNA-seq methods are prone to dropouts and relatively low-coverage of the data. We therefore cannot exclude that condition-specific programs would emerge in deeper or higher-coverage data.

While the eigenvector loadings do not clearly separate between disrupted and regulated, the magnitude of the corresponding eigenvalue is important to interpret how much a certain mode contributes to the overall correlation. We can score the genes with a weight that corresponds to their eigenvalue:

$$s_i = \frac{1}{P}\sum_{k=1}^{10}\lambda_k|v_i^{(k)}|, \tag{28}$$

where $P$ is the number of genes in the dataset, $v_i^{(k)}$ is the loading of gene $i$ in mode $k$ , and $\lambda_k$ is the eigenvalue of mode $k$ (sorted from highest to lowest). This score sums the eigenvector amplitudes, weighted by their corresponding eigenvalue, for the 10 highest modes (i.e. how much coordinated variance each gene carries). Now we can define the weighted number of participating genes as the number of genes that cross a given threshold. For any reasonable threshold we observe that regulated samples contain more 'correlated genes' than Dis-Arrest samples (for a threshold of $10^{-3}$: regulated median=231.5; Dis-Arrest median=96; Mann-Whitney U=49.5, p-value=0.015).

**Conclusion** The eigenvector analysis can reveal the dominant pathways that contribute to the correlation eigenvalues and can also tell us which pathways are anti-correlated within the culture. However, the eigenvectors alone show similar patterns in both regulated and disrupted conditions and therefore do not tell us whether a sample is regulated or not. The information about the strength of correlations within these pathways is carried by the associated eigenvalues, and not by the eigenvector loadings. The interpretation is that the genes that carry the regulated/dysregulated signal are similar, but the level of regulation is expressed in the magnitude of the eigenvalues, which is exactly the readout of GMP-Cor. Additionally, by analyzing the number of participating genes weighted by their eigenvalue, we observe that regulated samples contain a significantly larger amount of 'correlated genes' compared to Dis-Arrest samples for any reasonable threshold.

### 4 Summarizing GMP-Cor for all datasets

We classify the single cell experiments as follows:

- Experiment 1a contains 4 datasets of *E. coli* cells exposed to SHX, leading to a Dis-Arrest state.
- Experiment 1b ran in parallel to 1a and contains 4 datasets of VapC induced cells along with one exponential control.
- Experiment 2 was performed separately and contains 3 datasets of Reg-Arrest cells, and 2 datasets of Dis-Arrest cells.
- We included analyses of exponentially growing datasets from Ma et. el[2] and McNulty et al.[3].

Extended Data Fig. S3 summarizes the correlation eigenvalue distributions of all the datasets. To visualize the global correlativity of the data, 3 factors must be taken into account:

1. A low eigenvalue peak which is higher than the MP distribution and the scrambled data.

2. A heavy tail of eigenvalues that exceeds the largest eigenvalue of the MP distribution and the scrambled data. This feature is much better visualized in the CCDF plots of the distributions.

3. A large value of $\lambda_{max}$. Although this is not a robust, global feature, it gives a strong indication on the correlativity of the data.

While most datasets that exhibit correlations possess these 3 features, in some datasets only 2 are observed (for example, the exponential dataset from experiment 1b does not show a low eigenvalue peak, but the other 2 features do exist).

Calculating GMP-Cor for each dataset gives an indication of how correlative the data is. To test the correlation between this measure and the biological classification of the data, we divide all the datasets into two groups: disrupted and regulated (regulated includes also the exponential and 2-5h VapC datasets). We perform a Mann-Whitney U test on this classification, which returns a significant p-value (Fig. 3E).

### 5 Normalization of bulk RNA sequencing

Proper normalization of bulk RNA sequencing is crucial to achieve meaningful results for differential gene expression analysis. This is because differences in overall RNA counts often stem from technical variability, due to differences in library preparation, RNA extraction efficiency, sequencing depth, etc.[4]. *DESeq2*'s median-of-ratios normalization scheme, computes a scaling factor per sample based on the median of the ratios of each gene's count to the geometric mean across all samples, and normalizes the overall expression between conditions [5]. This method is mostly suited for cases in which we do not expect a large change in absolute RNA counts between conditions. When comparing growth-arrested to exponentially growing bacteria (that do not share a similar number of overall RNA counts) the resulting volcano plot is roughly symmetric - which can be misleading (the same amount of genes appear to be up and down-regulated, see Extended Data Fig. S12A,B). To test the validity of the GO enrichment results, that are based on the DEGs obtained from *DESeq2*, we also perform normalization by ERCC spike-ins. ERCC spike-ins are synthetic RNA molecules that are added to the sample at a known concentration. By analyzing the reads that map to these spike-ins we can normalize the data to account for technical variation only, without removing the overall expression difference that stems from biological variation. This is done by calculating the median of ratios between the spike-in counts to their known concentrations, and using this as a scaling factor for each sample. Following this normalization method, we clearly see an asymmetry between the up and down-regulated genes. This asymmetry is more pronounced in Reg-Arrest than in Dis-Arrest (Extended Data Fig. S12D,E). Nevertheless, GO enrichment analysis gives almost identical results for both normalization schemes, suggesting that the differences in overall transcripts between conditions does not affect our conclusions. The GO enrichment $p_{adj}$ values for Reg-Arrest and Dis-Arrest are presented for *DESeq2* normalization and ERCC normalization (Extended Data Fig. S12C/F respectively).

### 6 Validating GMP-Cor on mammalian cells

To test whether our framework for analyzing correlation patterns in scRNA-seq is applicable to other systems, we analyzed data on a cancer cell-line under treatment published by Oren et al.[6]. This work investigated persistence in a PC9 cancer cell-line treated with osimertinib. scRNA-seq was performed at multiple time points during treatment, and a fluorescent marker was tracked in order to sort cells into 3 groups after 14 days of treatment: proliferating, arrested and intermediate cells. We analyzed the scRNA-seq data of these 3 groups and asked whether GMP-Cor can distinguish between the level of correlations in these cells. We expected the proliferating group to show stronger correlations, because active growth requires the coordinated activity of many genes. The results show a clear reduction of GMP-Cor in the arrested compared to the proliferating group (Extended Data Fig. S15). GMP-Cor is elevated in all cell groups compared to the bacteria values, possibly reflecting the stronger coordination and expression levels in mammalian cells. It remains difficult to conclude whether the arrested PC9 cells are truly dysregulated, or just less tightly regulated than the proliferating cells, which would require further physiological measurements. Nevertheless, the results clearly show the expected trend, supporting our claim that GMP-Cor is a general scRNA-seq metric, and not restricted to the study of bacteria under stress.
