## Supplementary Figures for "GMP-Cor: Quantifying Transcriptional Dysregulation in Antibiotic Tolerance and Persistence"

### Extended Data Figures

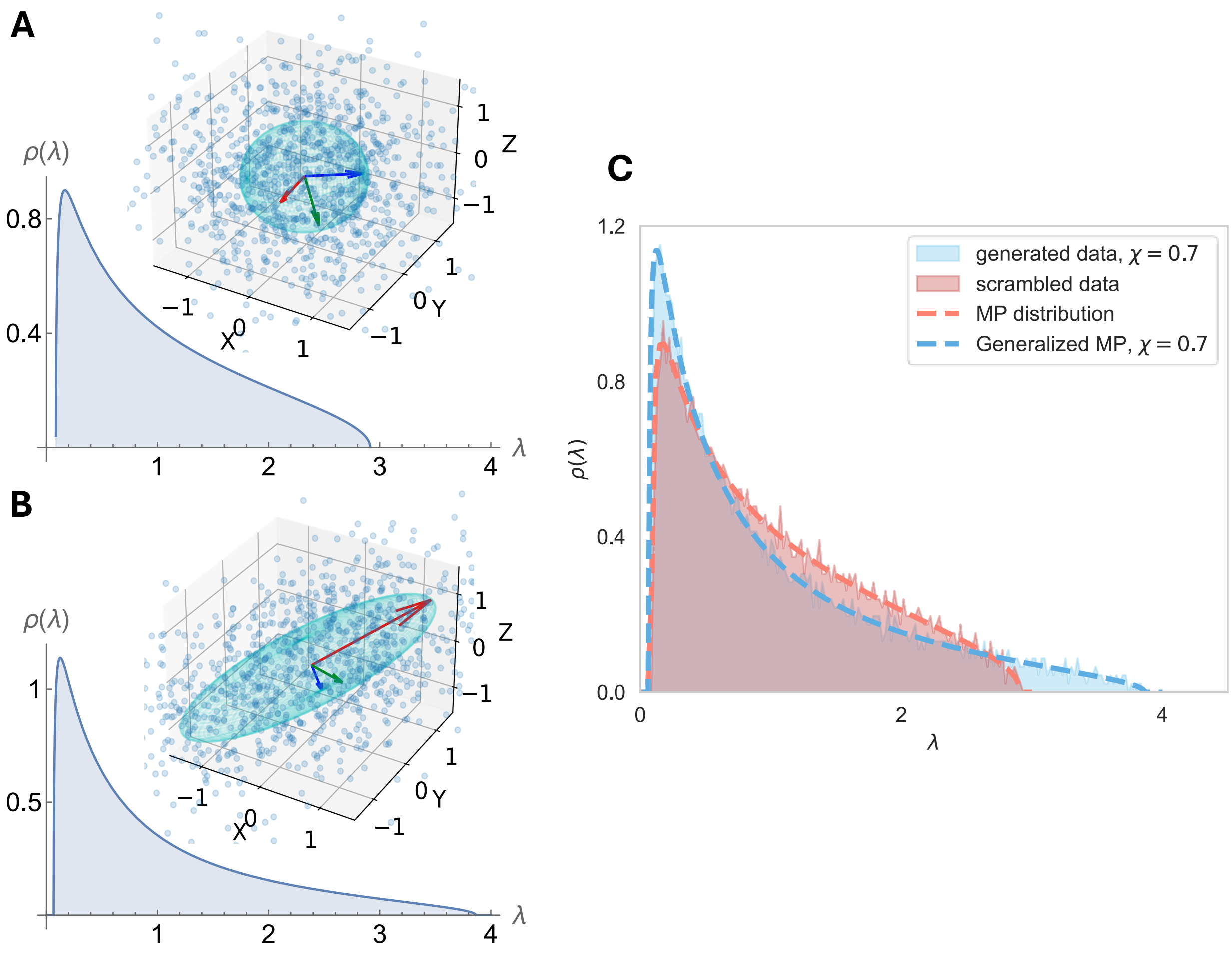


**Extended Data Figure S1: Visualizing the results of the Generalized MP (GMP) distribution**. (**A**) Analytical plot of the MP distribution. Inset: Scatter plot of identical and independently distributed (iid) 3-dimensional vectors, generated from a multivariate Gaussian ensemble. The sphere represents the scatter of the data, and is spanned by the eigenvectors of the correlation matrix (red, green and blue arrows). (**B**) Analytical plot of the non-iid generalization of the MP distribution. Inset: Same as in (A), but with variable dependencies. (**C**) The analytical results for $\alpha = 0.5$, $\chi = 0.7$ and $\alpha = 0.5$, $\chi = 0$ (dashed blue and red lines, respectively). Distribution of computer generated data with $\alpha = 0.5$, $\chi = 0.7$ is plotted in the shaded blue area, validating the analytical calculation. The data is then scrambled and its distribution is plotted in the shaded red area, coinciding with the original MP analytical result.

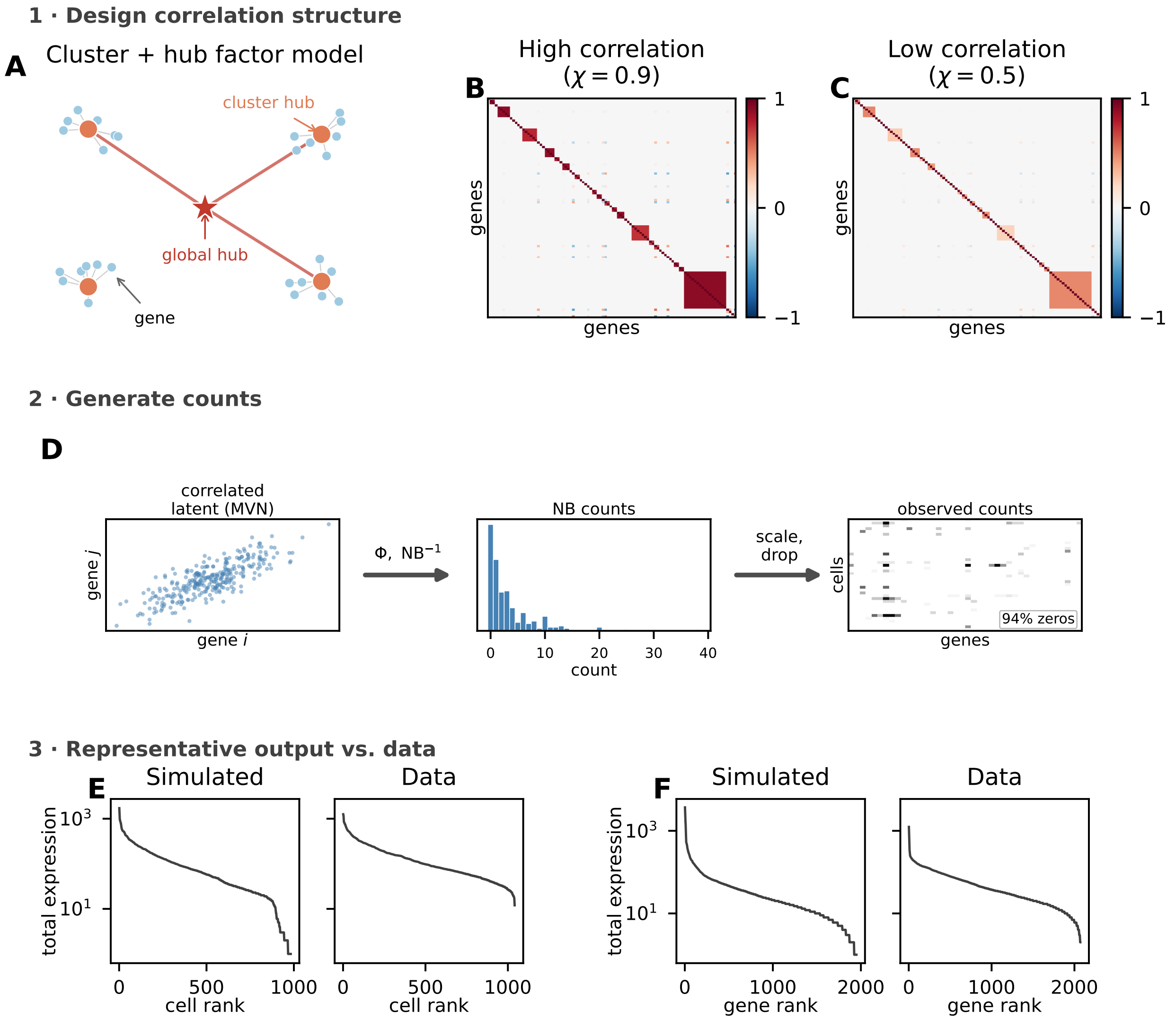


**Extended Data Figure S2: Design and validation of the synthetic single-cell RNA-seq data generator.** **(A)** Schematic of the cluster-hub model used to define the gene-gene correlation structure. Each gene (blue nodes) belongs to a cluster whose members are correlated through a cluster hub (orange nodes). A randomly selected subset of cluster hubs is additionally linked to a single global hub (red star), introducing long-range correlation between clusters. Edge weights are drawn from a uniform distribution and are modulated by the correlation strength parameter $\chi$. **(B, C)** Representative $100 \times 100$ sub-blocks of correlation matrices generated with high ($\chi = 0.9$, **B**) and low ($\chi = 0.5$, **C**) correlation strengths. Block-diagonal structure reflects the cluster topology. **(E)** Flow diagram of count generation via a Gaussian copula (link distribution). *Left to right:* (i) Correlated latent variables are drawn from a multivariate normal distribution $\mathcal{N}(\mathbf{0}, R)$, (ii) Latent variables are mapped to a unit interval via the Gaussian CDF ($\Phi$) and the negative-binomial quantile function is applied to each uniform variate; the per-gene mean $\mu_i$ is drawn from an inverse-Gamma distribution to reproduce the heavy-tailed expression distribution observed in real scRNA-seq data. (iii) Cell library-size variability is introduced by multiplicative log-normal amplitudes, followed by expression-dependent dropout ($P(\text{dropout}) = e^{-\lambda x}$) to generate the sparse observed count matrix. **(E, F)** Rank plots of per-cell (**E**) and per-gene (**F**) total counts on semi-log axes, confirming that the simulated data mimics the expression distributions characteristic of real single-cell data. The corresponding rank plots from a representative dataset are shown for comparison.

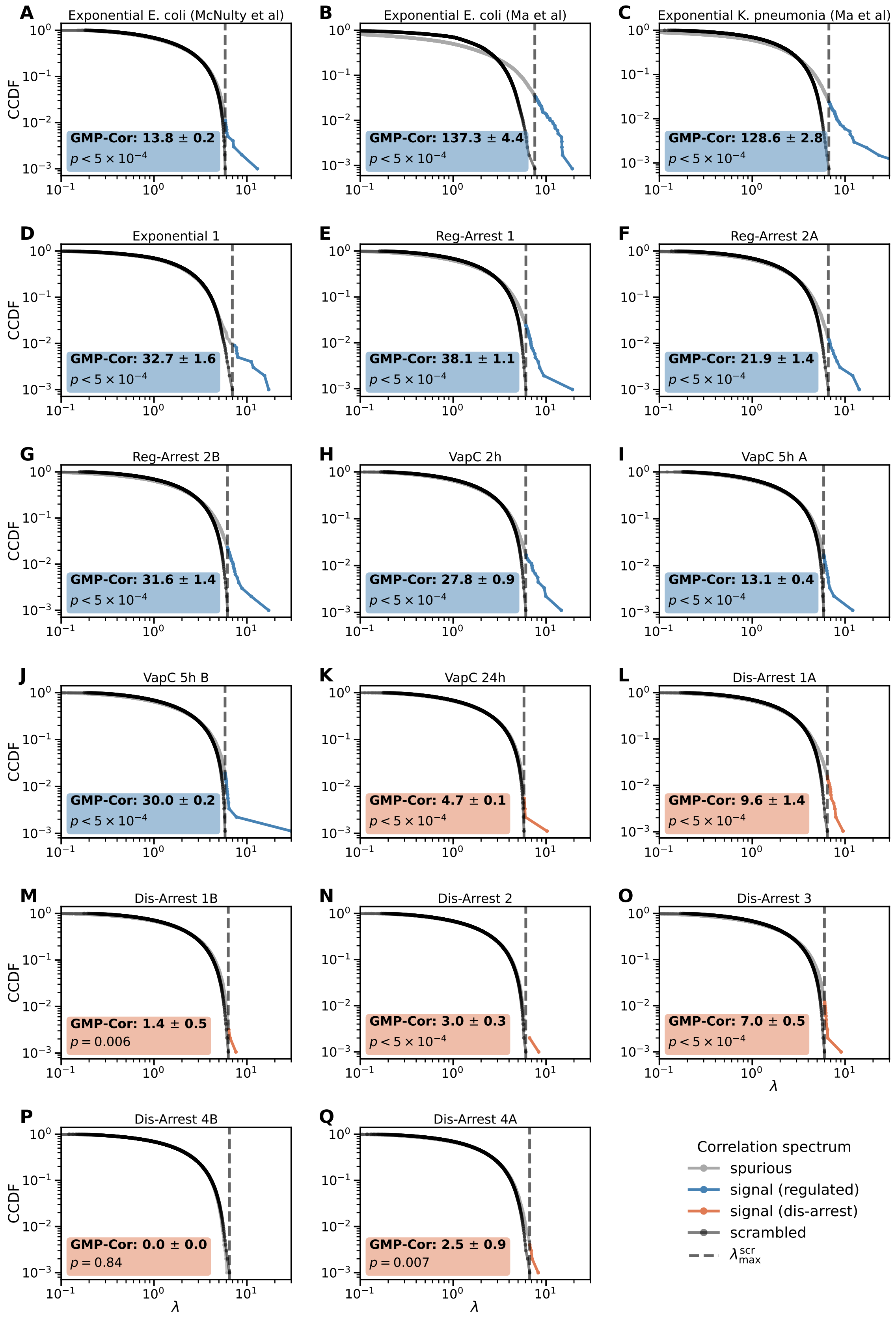


**Extended Data Figure S3: Overview of GMP-Cor of all datasets.** Log-log CCDF of correlation eigenvalues for all datasets analyzed. The dashed vertical line marks $\lambda_{max}^{scr}$ - eigenvalues exceeding this threshold (colored) reflect biologically meaningful gene-gene correlations absent in the scrambled control (black). Statistical significance of the original vs. scrambled distributions is obtained by a permutation test (methods) with the resulting p-value annotated in each plot. GMP-Cor value and confidence interval are annotated in each plot. Different numbers for the same condition represent biological replicates while different letters are technical replicates.

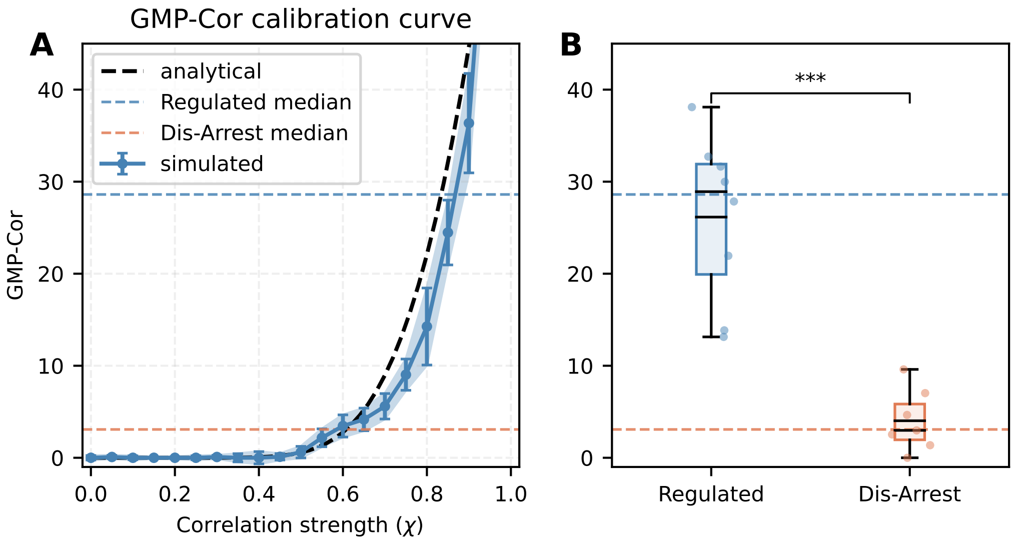


**Extended Data Figure S4: Calibration of GMP-Cor observable measure to the theoretical correlation strength**. **(A)** GMP-Cor calibration curve obtained by sweeping $\chi$ from 0 to 1 in the synthetic data generator model (n = 10 replicates per $\chi$ value), points show median ± s.d. Horizontal reference lines indicate the median GMP-Cor of the Regulated (blue) and Dis-Arrest (red) experimental groups, anchoring the expected GMP-Cor value of the two biological conditions to the correlation strength parameter derived from the numerical model. Dashed black line is the same calibration calculated from the analytical GMP distribution. **(B)** Distribution of GMP-Cor values across individual samples in the Regulated and Dis-Arrest groups (Replicated from Fig. 3E for easier comparison). Individual data points are overlaid on the box plots. Horizontal dashed lines represent the medians of each group. For comparison with the numerical model, y-axis is shared with panel (A). Statistical comparison by two-sided Mann-Whitney U test (***$p < 0.001$).

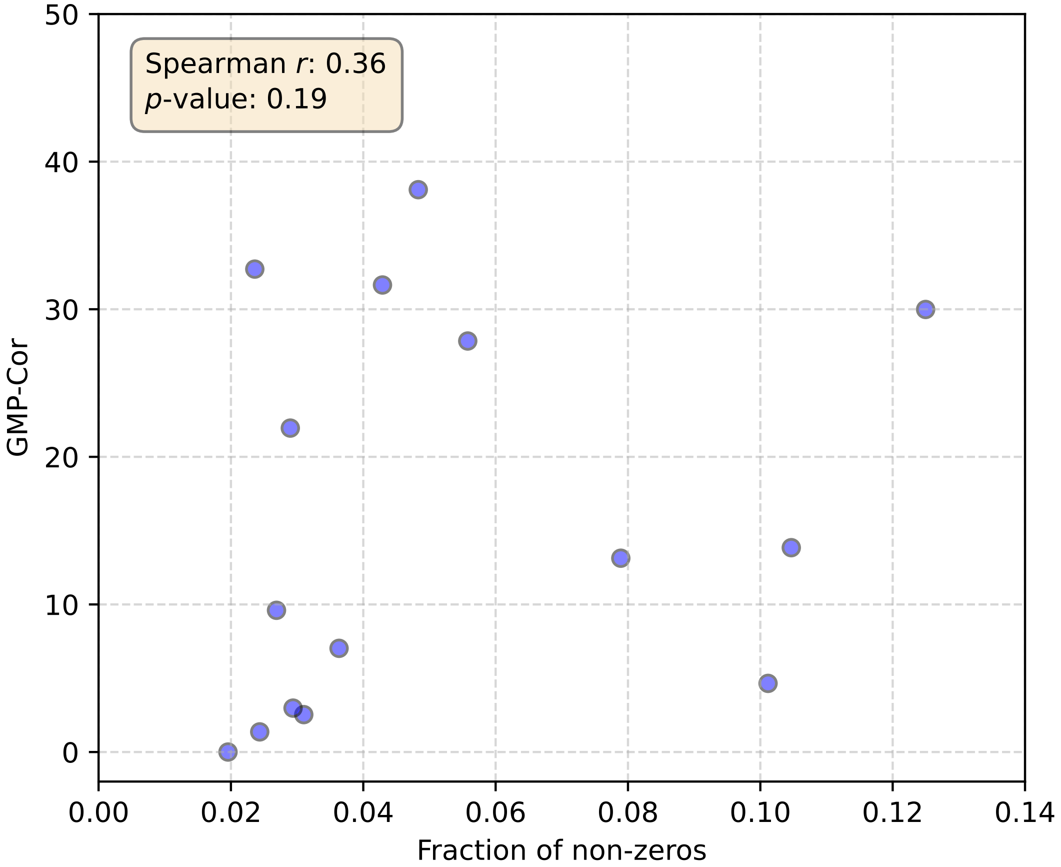


**Extended Data Figure S5: No significant correlation between sparsity and GMP-Cor metric**. Scatter plot of the GMP-Cor value against fraction of non-zeros for all datasets analyzed in this study. The resulting p-value of the Spearman-r correlation coefficient suggests that there is no correlation between these metrics, showing the validity of the GMP-Cor.

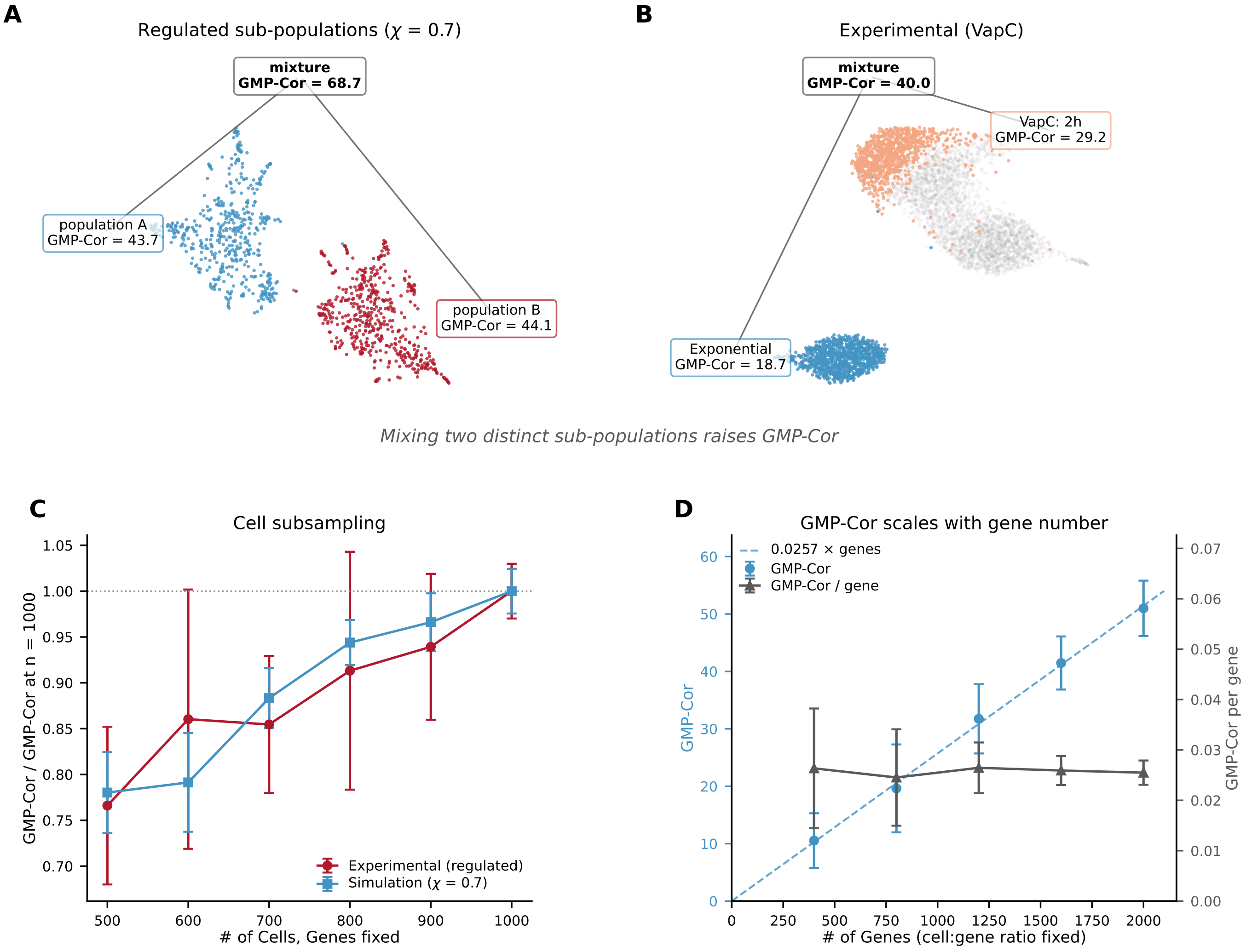


**Extended Data Figure S6: GMP-Cor is elevated by sub-populations and is mildly dependent on cell sub-sampling.** **(A-B)** UMAP embedding of distinct sub-populations with GMP-Cor calculated on each population and the mixture with a fixed cell/gene dimension. **(A)** Populations created by simulating data with high correlations ($\chi = 0.7$ - regulated). **(B)** Experimental sub-populations analyzed from the UMAP presented in Fig. 5C. Note that GMP-Cor does not exactly reproduce the calculated per sample GMP-Cor presented in Fig. 5G because it is calculated here on the highly variable genes of the union dataset, which do not necessarily capture the strongest correlation signal for each dataset separately. **(C)** GMP-Cor as a function of number of cells, randomly sub-sampled from the full dataset, for both simulated (blue) and a representative regulated dataset (red). **(D)** GMP-Cor as a function of gene number on simulated data shows a linear relationship, suggesting that GMP-Cor is an extensive quantity. Right axis: GMP-Cor normalized by gene number is flat over the presented range.

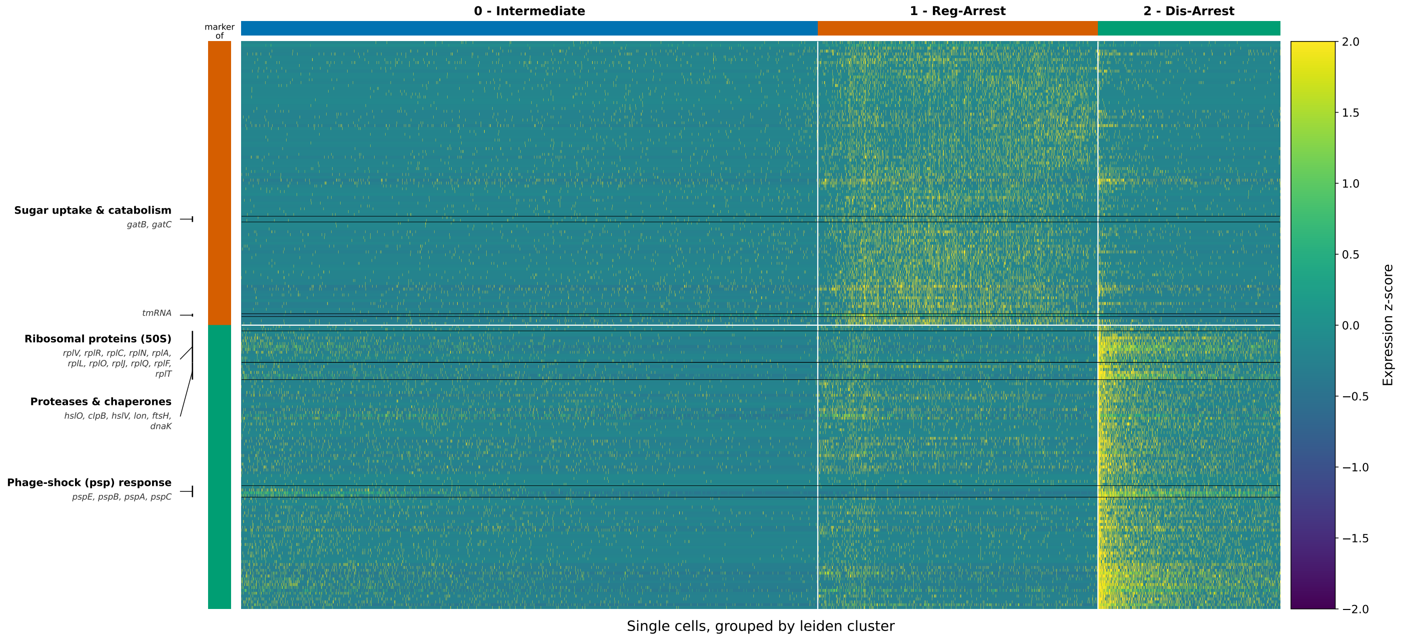


**Extended Data Figure S7: Heatmap of marker genes across single cells from SHX induced growth arrest and Reg-Arrest.** Columns are single cells grouped by Leiden clusters, rows are genes, with values showing per-gene z-scores across cells. Genes are ordered by hierarchical clustering on cell-level expression profiles, colored bars group marker genes with their associated leiden cluster. For orientation, gene groups that are discussed in the main text or strongly differentiate between the clusters are indicated on the left. Cluster composition is enriched as follows: cluster 0 - intermediate, cluster 1 - Reg-Arrest, cluster 2 - Dis-Arrest. For each Leiden cluster, the top 100 marker genes ranked by Wilcoxon test score are presented. Only marker genes with $p_{adj} < 0.05$ were kept for the analysis. Genes occurring in multiple clusters were de-duplicated.

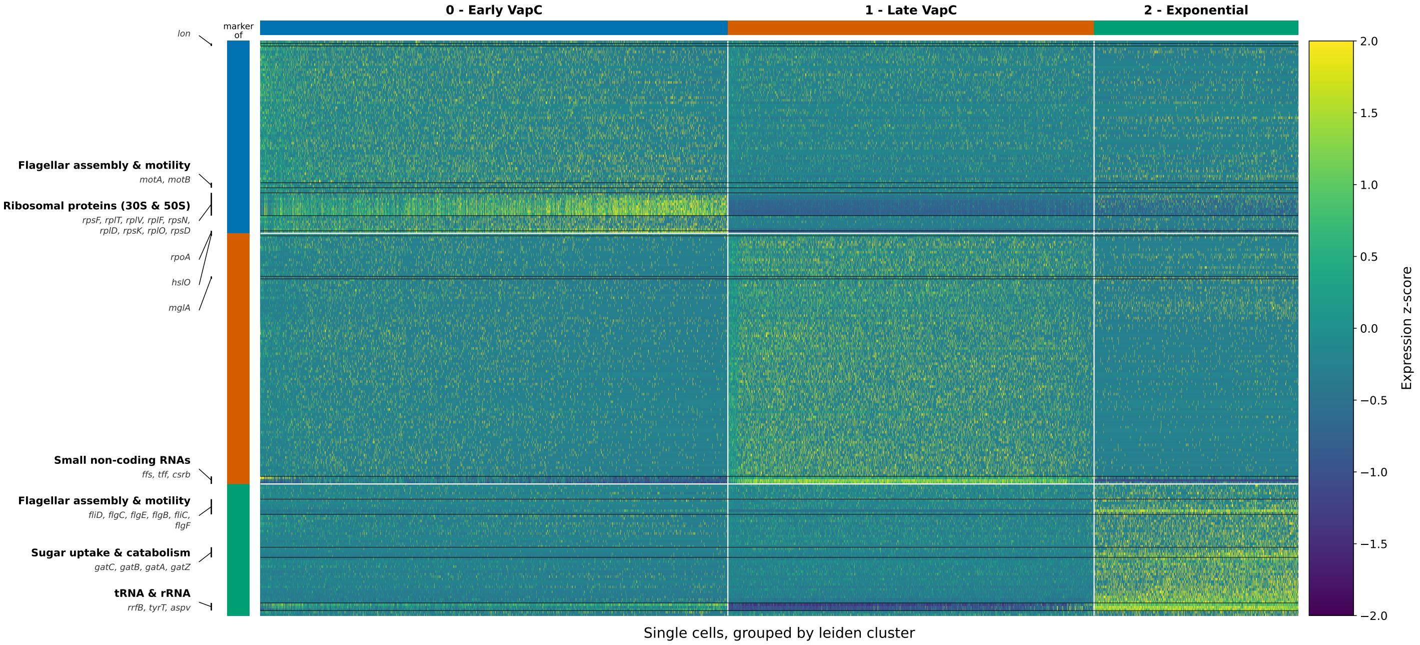


**Extended Data Figure S8: Heatmap of marker genes across single cells at different times in VapC induced growth arrest.** Columns are single cells grouped by Leiden clusters, rows are genes, with values showing per-gene z-scores across cells. Genes are ordered by hierarchical clustering on cell-level expression profiles, colored bars group marker genes with their associated leiden cluster. For orientation, gene groups that are discussed in the main text or strongly differentiate between the clusters are indicated on the left. Cluster composition is enriched as follows: cluster 0 - Early VapC induction (2-5 hours), cluster 1 - late VapC induction (5-24 hours), cluster 2 - Exponential (before induction). For each Leiden cluster, the top 100 marker genes ranked by Wilcoxon test score are presented. Only marker genes with $p_{adj} < 0.05$ were kept for the analysis. Genes occurring in multiple clusters were de-duplicated.

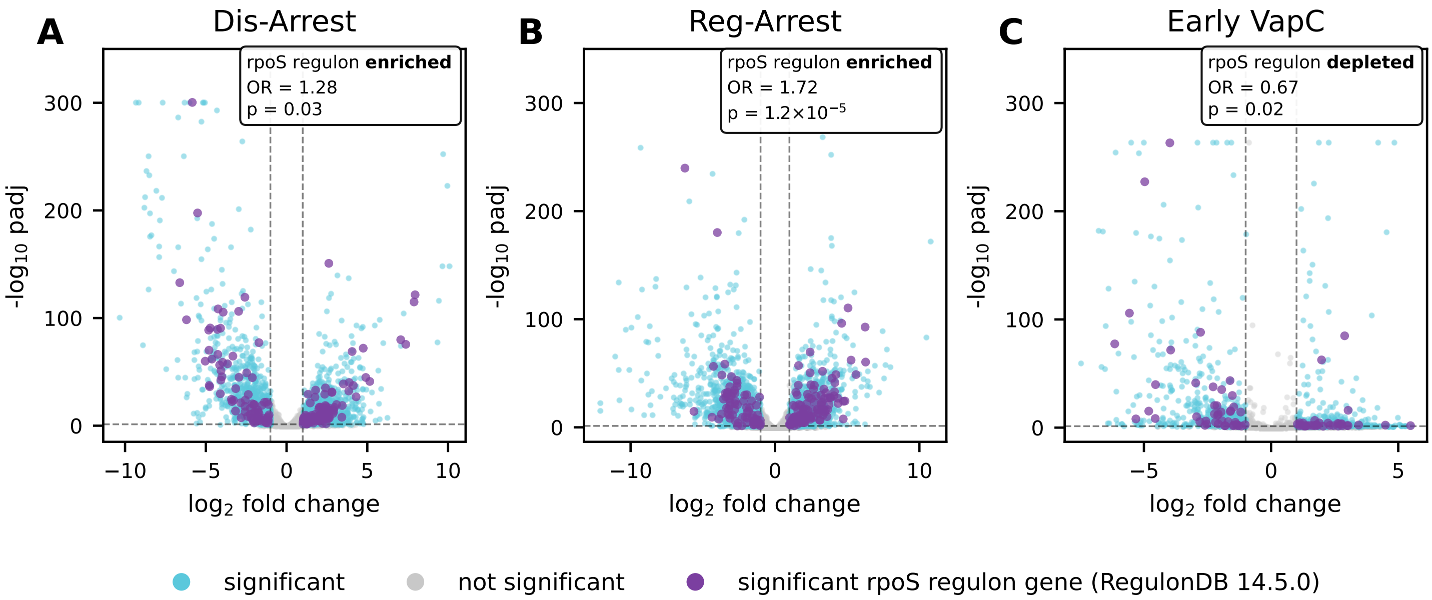


**Extended Data Figure S9: Expression of RpoS regulon genes in Dis-Arrest, Reg-Arrest and VapC compared to exponential growth.** **(A-C)** Volcano plots of differentially expressed genes (DEG) in Dis-Arrest **(A)**, Reg-Arrest **(B)** and 2 hours after VapC induction **(C)** compared to exponential growth. Significant genes ($|LFC| > 1$ and $p_{adj} < 0.05$) are marked in cyan, significant genes labeled as regulated by RpoS (according to RegulonDB 14.5.0) are marked in purple and non-significant genes are marked in grey. Enrichment test is performed in the up-regulated portion of the DEG's by Fisher's exact test. Odds ratio (OR), direction (enriched/depleted) and enrichment test p-values are stated in each plot. Both Dis-Arrest and Reg-Arrest show up-regulation of RpoS regulon (though the effect is much stronger in Reg-Arrest), while VapC shows a depletion of RpoS genes in the up-regulated gene set. (A-B) are based on bulk RNA-seq data, while (C) is performed on aggregated scRNA-seq data.

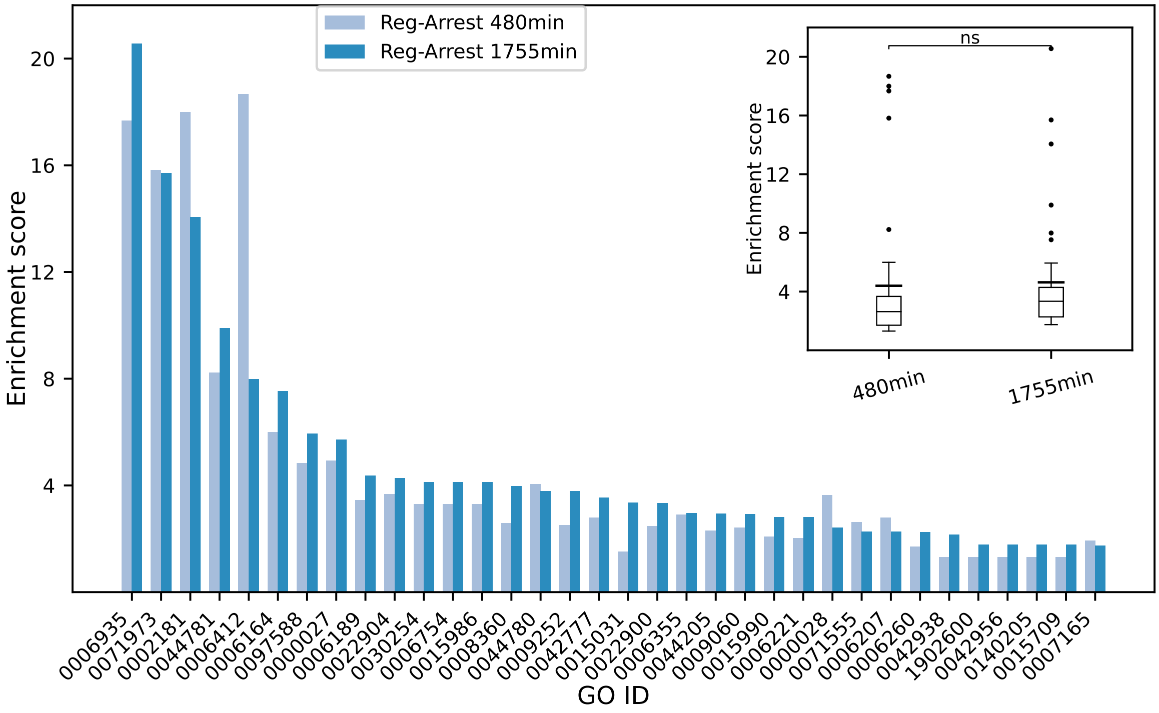


**Extended Data Figure S10: Comparison of GO term coherence between early and late time in Reg-Arrest.** Enrichment score ($-\log_{10}(p_{adj})$) of GO terms that are significantly enriched in both early (480min) and late (1755min) Reg-Arrest. Light blue represents the early time-point and dark blue represents the late time-point. Inset: Box plot of enrichment scores of all GO terms. The Mann-Whitney U-test was performed to show that there is no significant difference in the overall coherence of GO terms.

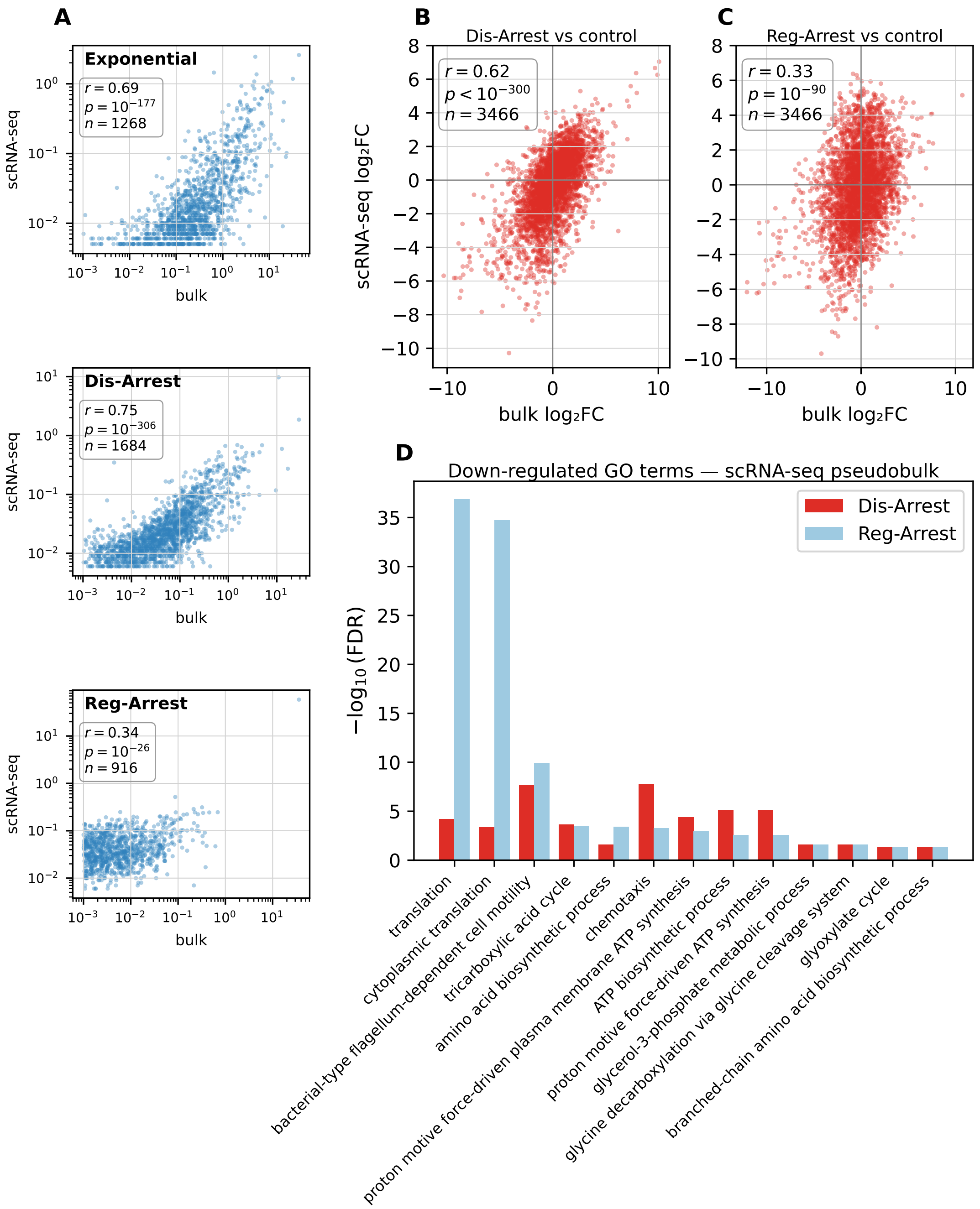


**Extended Data Figure S11: Correlation between scRNA-seq and bulk RNA-seq expression profiles.** **(A)** Per-gene mean expression in scRNA-seq versus bulk RNA-seq across three matched conditions (top: exponential, center: Dis-Arrest, bottom: Reg-Arrest), plotted on log-log axes. Each point represents one gene; only genes expressed above background in both datasets are shown (Pearson $r$, p-value and $n$ are indicated per panel). Pearson $r$ is computed on $\log_{10}$-transformed values. Bulk expression is normalized to molecules per cell units, achieved by ERCC normalization and cell count estimation. **(B, C)** Per-gene $\log_2$ fold-change (LFC) compared between the sc pseudobulk DESeq2 analysis and the bulk RNA-seq DESeq2 analysis, for Dis-Arrest **(B)** and Reg-Arrest **(C)** versus the exponential-phase control. The pseudobulk analysis aggregates per-condition single-cell counts into mean gene expression profiles and applies the same DESeq2 pipeline used for bulk data (Pearson $r$, p-value and $n$ are indicated per panel). **(D)** GO enrichment of down-regulated genes identified by the sc pseudobulk DESeq2 analysis (top 500 genes by LFC magnitude, matched study sets). Bars show $-\log_{10}(p_{adj})$ for GO terms that reach significance ($p_{adj} < 0.05$) in both Dis-Arrest (red) and Reg-Arrest (blue). The same functional categories suppressed in the bulk analysis (translation, flagellar motility, chemotaxis, energy metabolism) are independently recovered in the single-cell data, validating the biological signal detected across both platforms. Reg-Arrest displays stronger enrichment than Dis-Arrest for most terms, consistent with the bulk GO results in Fig. 4B.

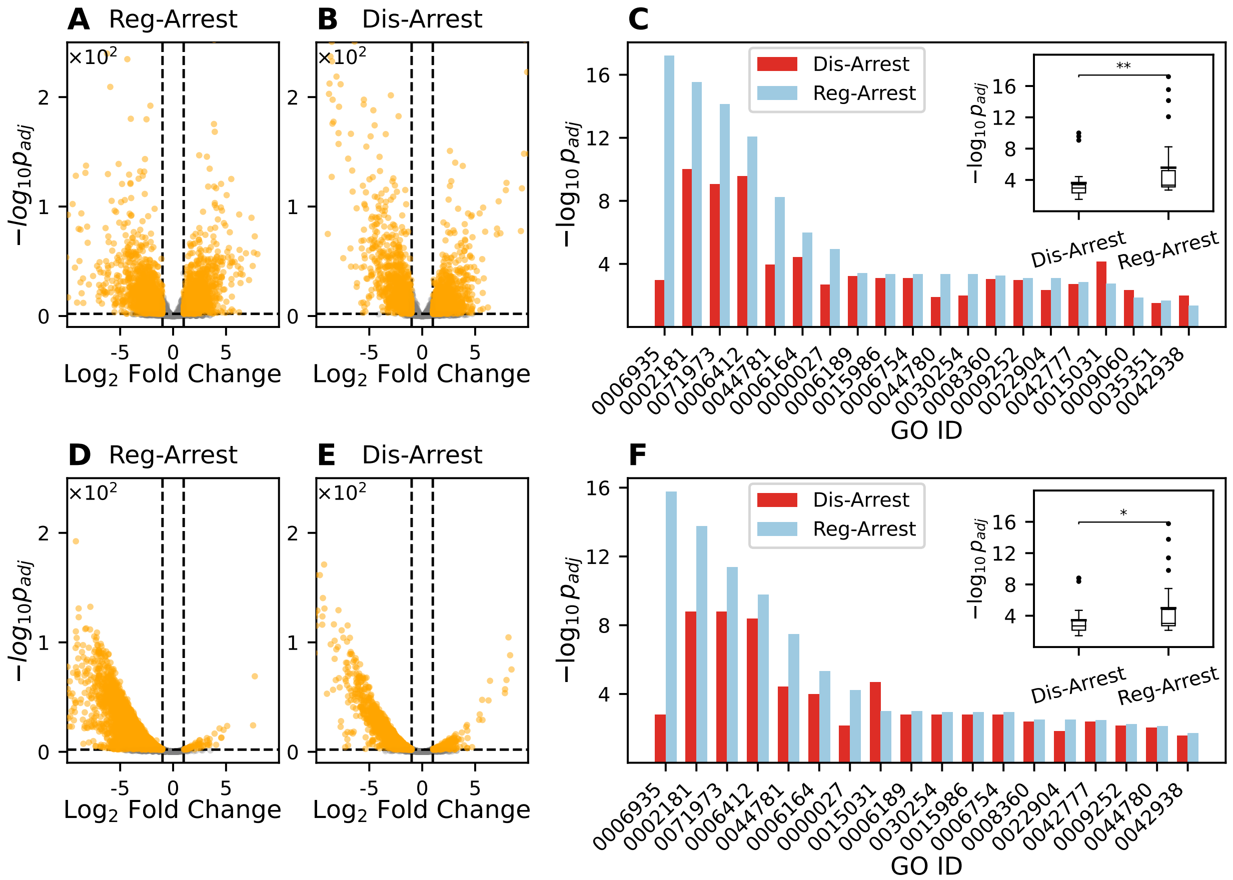


**Extended Data Figure S12: Comparison of normalization methods for bulk RNA sequencing. (A-B)** Volcano plots of the bulk RNA expression compared to exponential cells using the *DESeq2* normalization scheme. **(D-E)** Volcano plot of the bulk RNA expression compared to exponential cells using ERCC spike in normalization. Scaling factors for each sample were obtained by calculating the median ratios of ERCC spike ins (Methods) and then differential expression analysis was performed using the *DESeq2* algorithm. **(C,F)** Adjusted p-values from GO enrichment analysis, using the DEGs obtained from *DESeq2* normalization (C) and ERCC normalization (F). Similar results are obtained by both methods.

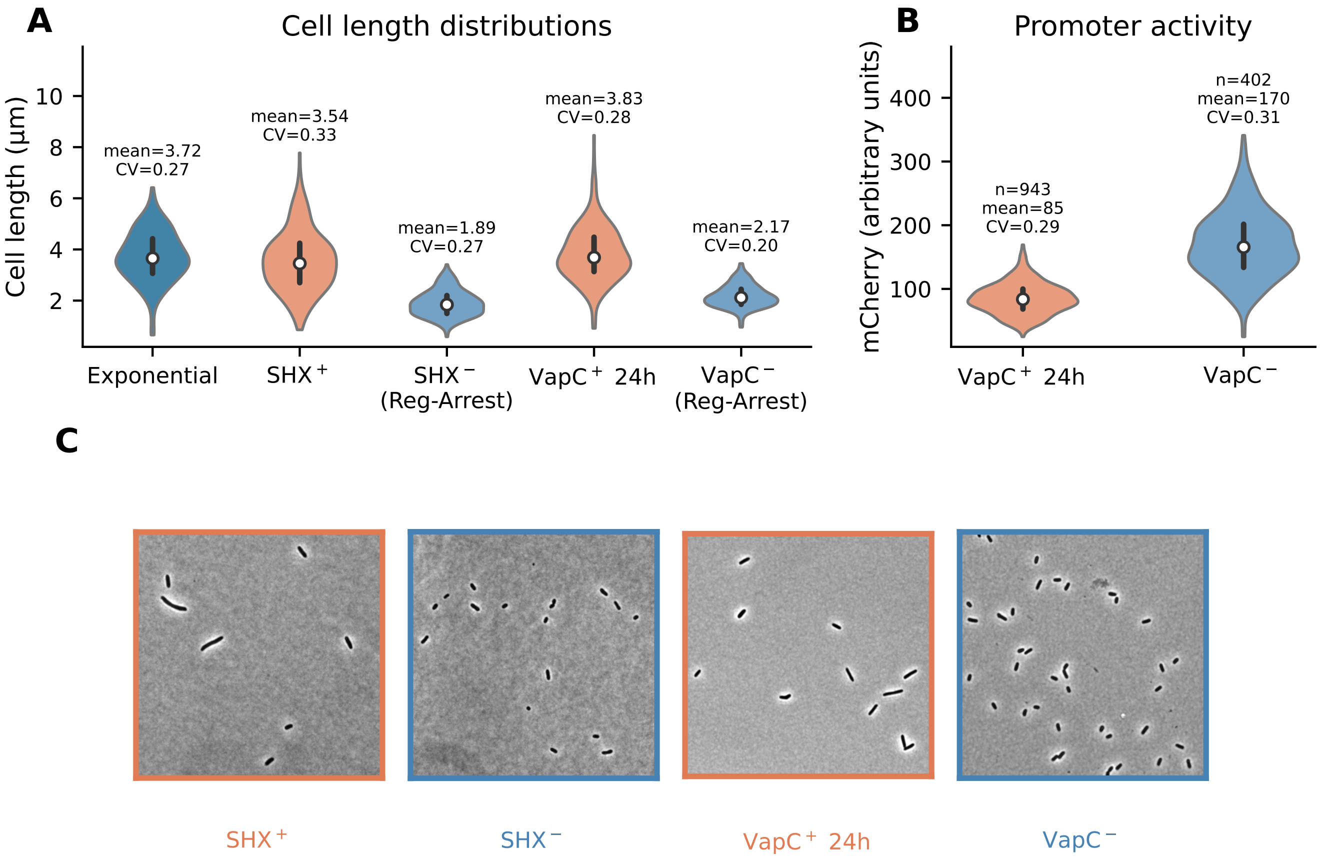


**Extended Data Figure S13: Single-cell morphology and expression levels for SHX and VapC conditions against Reg-Arrest control.** **(A)** Cell length distributions analyzed from microscopy images of different conditions. Mean and CV are shown above each distribution. Blue/red distributions represent Regulated/Dis-Arrest conditions respectively. Distributions are sub-sampled to match the sample size of the distribution with the minimal number of cells ($n = 233$). **(B)** Single-cell fluorescent intensity distribution of an induced marker (mCherry) attached to the same promoter as *vapC* for the VapC$^+$ and VapC$^-$ 24h condition. The variability in mCherry fluorescence corresponds to the variability in VapC expression. CV of VapC production is comparable to the promoter activity of the VapC$^-$ strain. **(C)** Representative microscopy images of the different experimental conditions, highlighting the elongated, larger area in the Dis-Arrest conditions (Red frame) relative to Reg-Arrest (Blue frame).

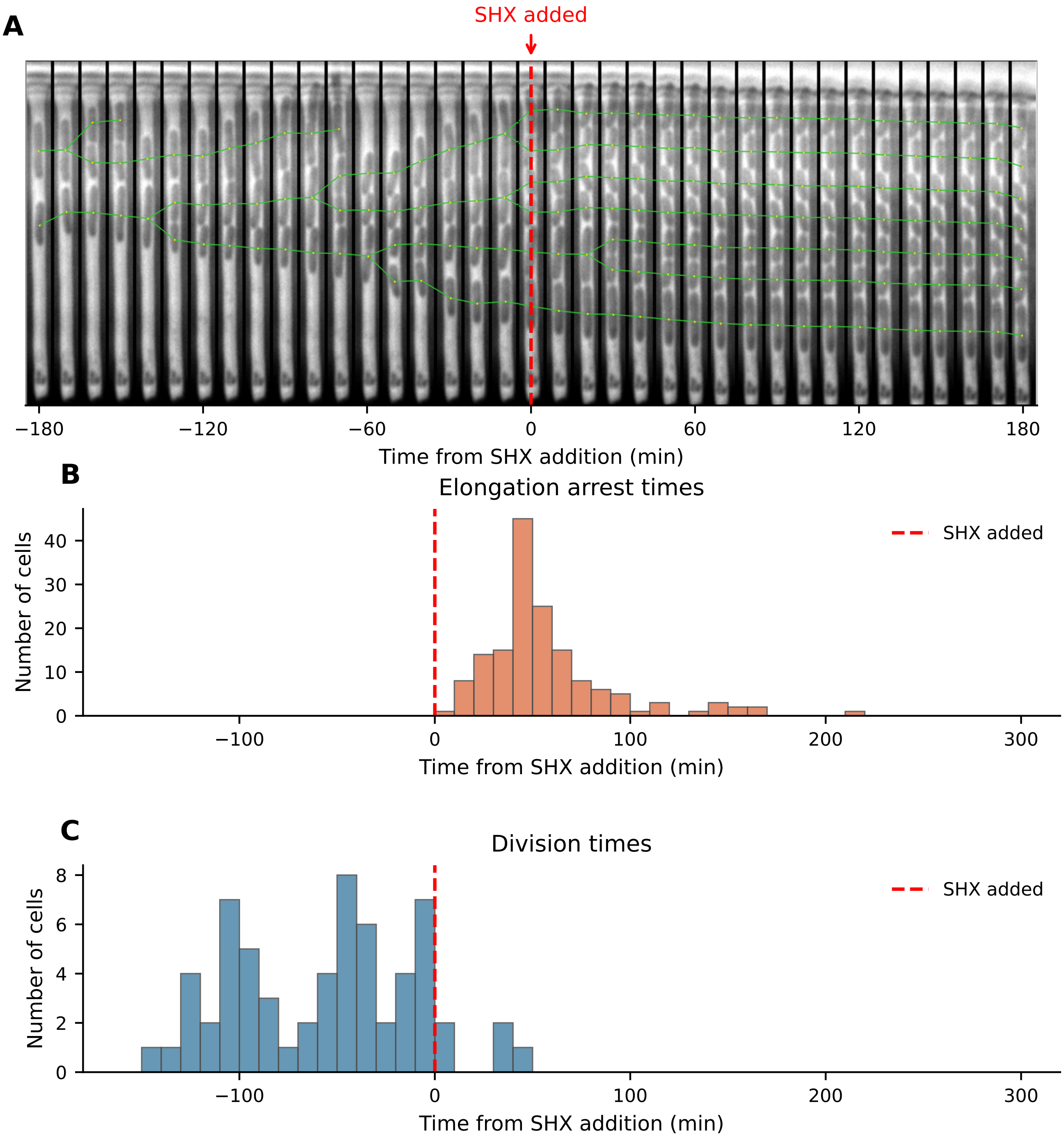


**Extended Data Figure S14: Growth-arrest dynamics after SHX exposure.** **(A)** Kymograph image of cell growth in a representative mother-machine trench. At $t = 0$min, SHX is added (red dashed line) and cells abruptly arrest growth. Green lines track lineages and divisions. **(B)** Distribution of times when single-cells stopped elongating after exposure to SHX ($n = 155$, median=40min, SD=66min). **(C)** Distribution of division times. Of all $n = 234$ evaluable cell tracks, 155 halted within the analysis window, 62 divisions were recorded (all up to and within 40 minutes of SHX addition), and 17 were ambiguous (did not divide but did not pass the criterion for elongation arrest). No track showed clear growth or division in the analysis window after $t = 40$min.

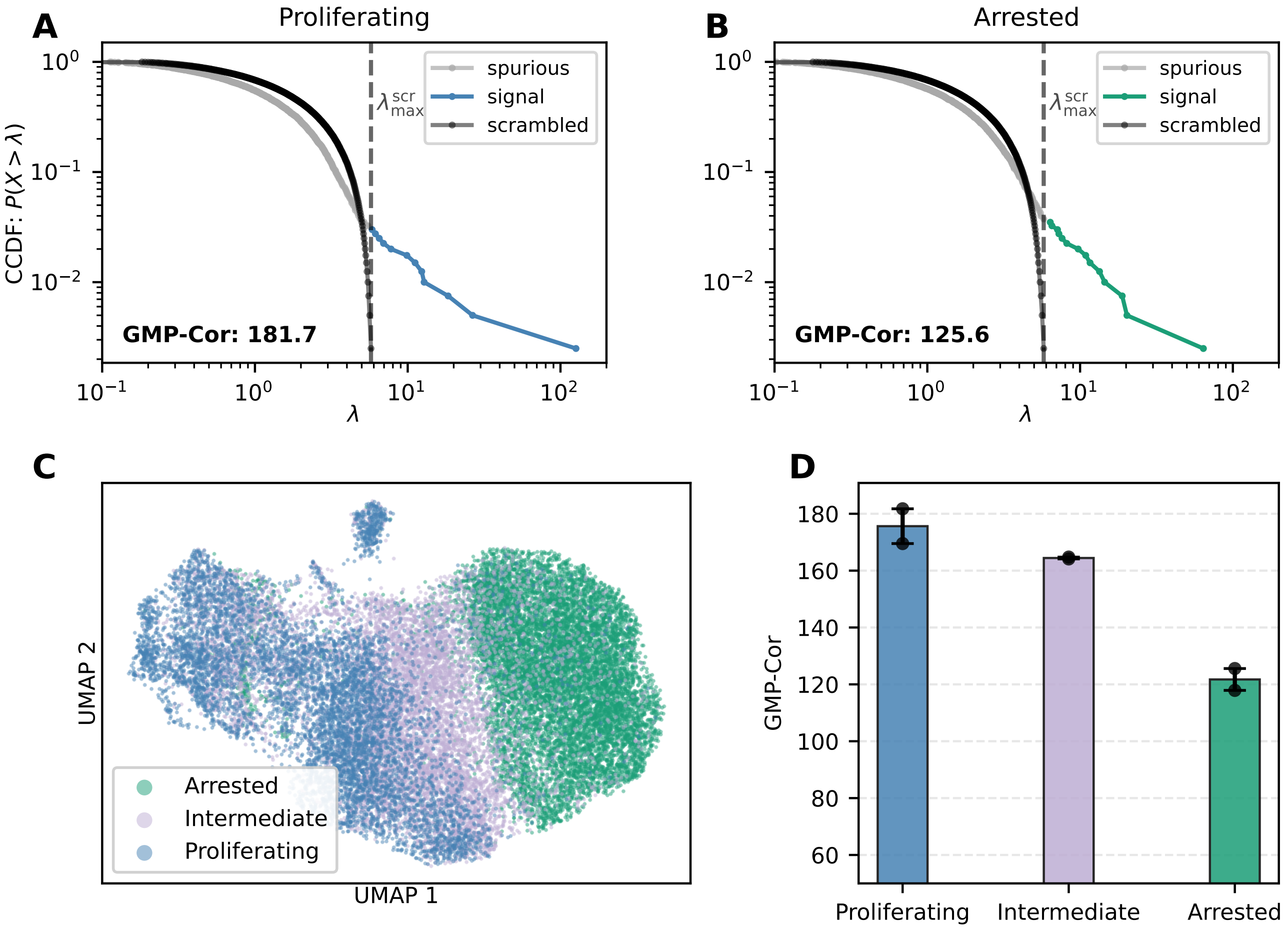


**Extended Data Figure S15: PC9 cancer cell line analysis of data obtained from Oren et. al.** [**6**]. **(A-B)** Eigenvalue distributions in log-log ccdf plots for both proliferating (A) and arrested (B) cells after 14 days of osimertinib treatment. The proliferating cells show a longer tail of eigenvalues and an elevated GMP-Cor compared to the arrested cells, suggesting stronger gene-gene correlations. **(C)** UMAP of all cells after 14 days of osimertinib treatment, the UMAP separates between the physiologically distinct growing and non-growing cells. **(D)** Bar plot summarizing GMP-Cor for all the samples in this dataset. The average GMP-Cor for each condition is shown as the level of each bar, with the error bars representing the standard error. The separate data-points are marked with black dots.

**A**

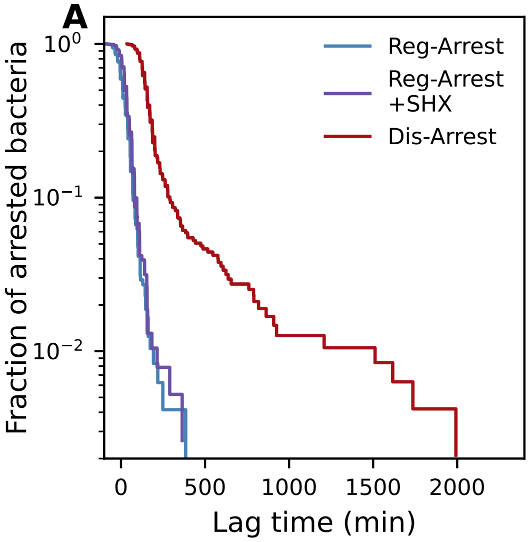


**B**

| Stress | Strain | Study | Lag time mean (min) | Std (min) |
|---|---|---|---|---|
| Reg-Arrest rep1 | KLYR | This study | 5 | 66 |
| Reg-Arrest rep2 | KLYR | This study | -14 | 72 |
| Reg-Arrest rep3 | KLYR | This study | 11 | 69 |
| SHX rep1 | KLYR | This study | 191 | 300 |
| SHX rep2 | KLYR | This study | 143 | 118 |
| SHX rep3 | KLYR | This study | 111 | 126 |
| Reg-Arrest+aTc | MG1655 | This study | 61 | 26 |
| VapC 4h | MG1655 | This study | 331 | 60 |
| VapC 24h | MG1655 | This study | 399 | 71 |
| SHX | KLYR | Kaplan et. al | 203 | 299 |
| Reg-Arrest | KLYR | Kaplan et. al | 34 | 47 |
| Reg-Arrest+SHX | KLYR | Kaplan et. al | 19 | 49 |

**Extended Data Figure S16: Summary of lag time distributions. (A)** CCDF plots of lag time distributions for Reg-Arrest, Reg-Arrest with SHX added, and Dis-Arrest. Lag time distribution of Reg-Arrest is not affected by adding SHX after entry into stationary phase. **(B)** Summary of lag time statistics for all conditions investigated in this study. Additional data is adapted from Kaplan et. al[1] to show that long and variable lag times - the hallmark of Dis-Arrest, are obtainable with many different types of stress and bacterial strains. Different Dis-Arrest conditions are in red while Reg-Arrest conditions are in blue.
